# Design of highly potent anti-biofilm, antimicrobial peptides using explainable artificial intelligence

**DOI:** 10.1101/2024.11.17.622654

**Authors:** Karina Pikalyova, Tagir Akhmetshin, Alexey Orlov, Evan F. Haney, Noushin Akhoundsadegh, Jiaying You, Robert E. W. Hancock, Dragos Horvath, Gilles Marcou, Artem Cherkasov, Alexandre Varnek

## Abstract

Antimicrobial peptides have emerged as a potential alternative to traditional small molecule antibiotics. They possess broad-spectrum efficacy and increasingly confront the challenges of bacterial resistance, especially the adaptive resistance of biofilms. However, advanced rational peptide design methods are still required to ensure optimal property profiles of such peptides, while limiting the cost of their synthesis and screening. Here we present a computational pipeline for the rational de novo design of antimicrobial and anti-biofilm peptides based on an explainable artificial intelligence (XAI) framework. The developed framework combines a Wasserstein Autoencoder (WAE) and a non-linear dimensionality reduction method termed generative topographic mapping (GTM). The WAE was used to learn the latent representation of the peptide space, while the GTM guided the generation of novel AMPs through an illustrative depiction of the latent space in the form of 2D maps. The efficacy of the peptides generated with the developed pipeline was experimentally verified by synthesis and testing for activity against methicillin resistant *Staphylococcus aureus* (MRSA), achieving a 100% hit rate in targeting biofilms. Notably, the most potent anti-biofilm peptide developed in this study demonstrated almost one order of magnitude improvement in IC_50_ value compared with the potent anti-biofilm peptide reference “1018”, used as a positive control. The developed pipeline is readily extendable for the optimization of additional peptide properties, including cytotoxicity, tendency to aggregate and proteolytic stability, underscoring its potential utility for rational design of the peptide-based therapeutics.

## Introduction

The advent of antibiotic medicines^1^ and their consistent and widespread use has been pivotal for treating infections ranging from mild to severe.^2^ However, this has led to bacterial adaptation and the emergence of antimicrobial resistance (AMR), exacerbated by antibiotic misuse and poor antibiotic stewardship, resulting in AMR becoming a critical public health challenge. The alarming rate of multidrug resistance (MDR), exemplified by pathogens such as methicillin-resistant *Staphylococcus aureus* (MRSA) and multidrug-resistant *Pseudomonas aeruginosa*, continues to limit treatment options and undermine effective infection management.^3^ An additional concern involves the fact that all currently-available antibiotics have been developed by targeting planktonic (free-floating) pathogens, as opposed to surface-associated bacterial biofilms which are more common (65% of all infections) and are substantially more resistant^4–6^, due to their alternative stress-adapted lifestyle, and their eradication is minimal at conventional antibiotic dosages^7^. Thus, there is an urgent need for innovative therapeutic strategies to circumvent planktonic- and biofilm-associated bacterial infections and improve treatment outcomes.

Antimicrobial peptides (AMPs) represent a prospective alternative to conventional small-molecule antibiotics, while the subset termed anti-biofilm peptides (ABP) represent a new agent effective against bacterial biofilms^8^. They possess advantageous properties such as a slow rate of evolved bacteria resistance^9^ and broad-spectrum activity^10^ often acting against both Gram-negative and Gram-positive bacteria.^11^ In nature, these peptides, collectively termed host defence peptides, are produced by multicellular organisms as a part of the innate defence system^12^. Host defence peptides have one or more of antibacterial, anti-biofilm or immunomodulatory functions^13^. They are characterized by their short length ranging from 10 to 50 amino acids, net positive charge^11^ varying between +2 and +9, presence of hydrophobic and cationic amino acids organized in way allowing them to exert amphipathic character and varying secondary structures^12^. Their cationic, amphipathic nature allows them to target negatively charged bacterial membranes, and undergo conformational transitions.^12,14,15^ The advantage of targeting biofilms includes the lack of FDA-approved drugs targeting biofilms and the opportunity to utilize topical therapy (reduced cost of goods and toxicity issues) since biofilms form on accessible surfaces.

While such peptides represent promising alternatives to antibiotics, particularly in targeting biofilms, rational drug design remains essential. In this regard, computational methods, including various Quantitative Structure-Activity Relationship (QSAR) models^3^, have been widely used. Das et al.^16^, for instance, suggested a framework for attribute-conditioned sampling of AMPs with the generative part based on a Wasserstein Autoencoder followed by long short-term memory (LSTM)-based classification of sampled peptides by properties and molecular dynamics simulations. This proved to be effective in accelerated generation of AMPs with experimentally verified broad-spectrum activity. Capecchi et al.^17^ designed short non-hemolytic AMPs using recurrent neural networks (RNN) fine-tuned through a transfer learning approach, using RNN classifiers and filters to enable potent AMP *in silico* screening, with experimental validation. Platforms similar to those described above have already shown effectiveness in the domain of small molecule discovery and are widely adopted in industry, as exemplified by Chemistry42 developed by Insilico Medicine^18^ and open-source REINVENT4 framework from AstraZeneca^19^.

Here, a novel pipeline for *de novo* design of anti-biofilm and antimicrobial peptides was based on the Wasserstein Autoencoder (WAE) coupled with the Generative Topographic Mapping (GTM). This framework, inspired by the methodology of Sattarov et al.^20^, is characterized by its ability to provide an illustrative visualization of latent space in the form of a 2D map (Fig. 1). This visualization facilitates the interpretability of complex data sets and allows rational analogue generation, aligning the framework with principles of explainable artificial intelligence (XAI) for drug design applications^21^, namely, transparency, justification, informativeness and uncertainty estimation.

**Figure 1.**
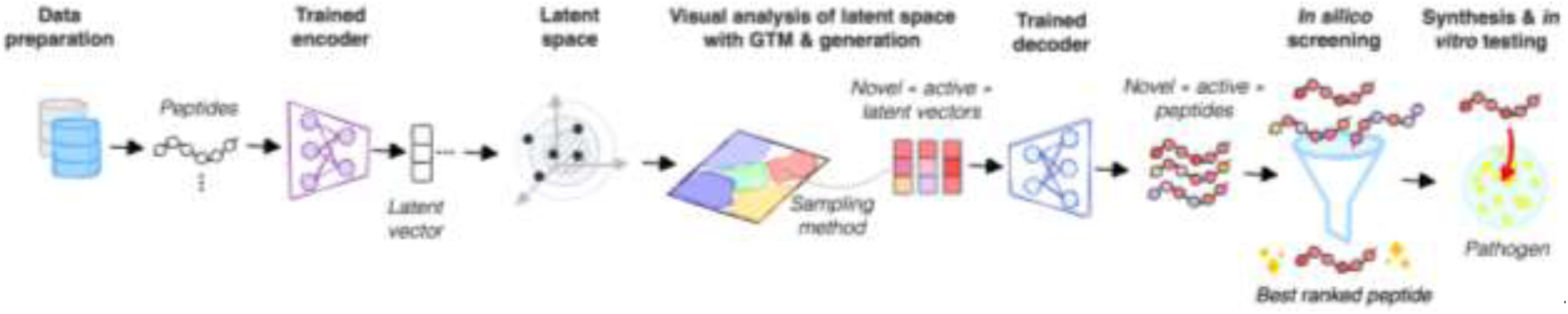
Overview of the pipeline developed in this work and used for successful generation of ABPs and AMPs.

The pipeline starts with data preparation encompassing data collection and pre-processing of peptide sequences. It is followed by the training of the Wasserstein Autoencoder (WAE) and encoding of the peptide sequences as dense vectors of the latent space. To visualize the latter in 2D, GTM is applied, enabling analysis of the characteristics of the distribution of the peptides in the latent space by the means of maps. This approach aided in navigating the latent space and identifying “landmarks”, representing zones populated by peptides with desired properties. These zones served as seeds to sample novel latent vectors, which were subsequently translated into peptide sequences with a decoder. The generated peptides were subjected to *in silico* screening based on machine learning models, resulting in a small set of eight peptides for experimental testing. When compared to the reference broad spectrum anti-biofilm peptide 1018^22,23^, all tested peptides were more active against MRSA biofilms and five out of eight peptides exhibited higher activity against planktonic cells. The experimental validation of the generated peptides, confirming their inhibitory activity and broad-spectrum character, demonstrated that the introduced framework effectively streamlines *de novo* generation of AMP hits. This approach holds potential for further user-defined multi-property-constrained *de novo* design, leveraging moderately sized datasets (∼45K).

## Results

### Data curation

To enable WAE learning, a dataset of 45,072 peptide sequences of lengths 10-14 composed of proteinogenic amino acids was collected. The dataset included 680 peptides from an experimentally validated in-house collection further referred to as “in-house” and other peptides retrieved from publicly available databases, namely TrEMBL^24^, DBAASP^25^, SATPdb^26^, SwissProt^24^, FermFooDb^27^, Hemolytik^28^, NeuroPedia^29^, BaAMPs^30^ and APD3^31^ (for more details see “Data collection for WAE training” subsection in methods and Table S1). The in-house peptides were labelled as active or inactive following the defined thresholds discussed in detail in the “Experimental dataset” subsection of methods.

### Wasserstein Autoencoder

The WAE was trained on the above-collected 45,072 peptides. The reconstruction rate was 98.7% on the validation set after 1000 epochs, with each epoch lasting 5 seconds. A detailed explanation of the WAE architecture is given in the corresponding Supplementary note in SI. The 256- dimensional mean latent vectors representing the centers of the peptide distributions served as vectors on which to build the GTM.

### GTM-driven analysis of the latent space

The GTM built on WAE latent vectors provides a human-readable representation of the latent space of peptides in the form of a 2D map through dimensionality reduction. The latent space data points representing peptides are fuzzily assigned to the nodes of the map. In greater detail, the position of a peptide on the 2D map is defined by a so-called responsibility vector, whereby the individual values reflect strength of association of the peptide to each node of the map. Thus, an individual peptide can be fuzzily represented by its responsibility “cloud” on the map, or, alternatively, by the dot corresponding to the barycenter of this responsibility cloud. Both fuzzy and barycentric representations will be used in the following. Responsibility vectors support the creation of fuzzy GTM landscapes colored according to either average values of peptide properties (activity, cytotoxicity, etc.) or by the probability to find sequences of a certain class (e.g., active/inactive) residing in a given node. To provide a general overview of the peptide motifs dominating in particular zones of the map, the density landscape was used (Fig. 2 center – darker nodes containing more peptides). The maps allowed the examination of peptide motifs shared by sequences residing in specific zones via sequence logo representations ^32^. For instance, the motifs of the peptides residing in the zones populated by active in-house peptides are shown in Fig. 2. They highlight alternating patterns of hydrophobic and hydrophilic residues within the peptides. Zone A in the density landscape grouped peptides that are rich in arginine and multiple hydrophobic residues such as tryptophan and isoleucine, featuring repeating RW units known to be effective antibacterial motifs/ pharmacophore^15,33^. Zone B was populated by peptides with both arginine and lysine cationic residues and primarily leucine and isoleucine residues, showcasing several triplet motifs such as MRR and RKI with a ratio of hydrophobic to hydrophilic residues of 1:2. The GTM-driven analysis of the zones hence allowed for the identification of the motifs inherent to AMPs and enhanced understanding of the sequence-activity relationships.

**Figure 2.**
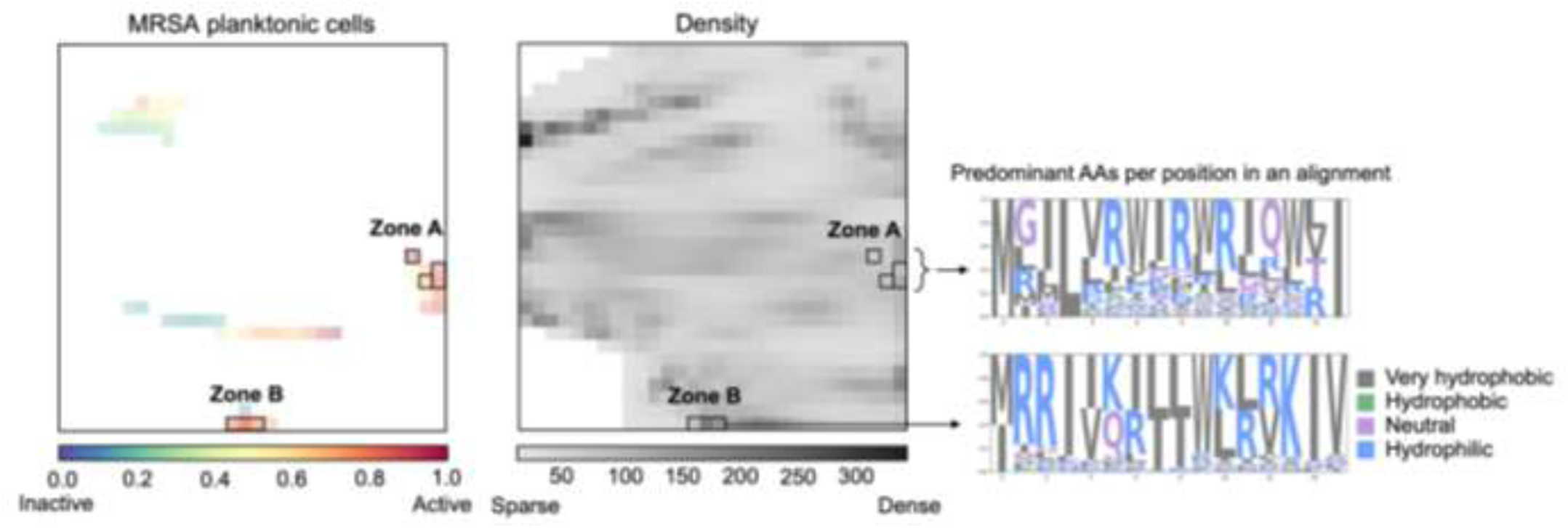
Left: Activity class landscape colored by the probability that active class in-house peptides (inhibiting MRSA planktonic cells) reside in the corresponding node, see Supplementary Fig. S1 A for the anti-MRSA biofilm activity class landscape. The intensity of color reflects the cumulated responsibility, i.e., fuzzy count of resident in-house peptides. Middle: Density landscape showcasing nodes densely populated by active peptides in black rectangles. The white zones correspond to empty (unpopulated) regions given a current dataset. Only nodes with a density higher or equal to 0.1 are shown on the density landscape. Right: Sequence motif maps of the peptides in active zones A and B.

The GTM also allowed for direct comparison of peptide datasets on the same map. For example, the comparative landscape delineating the zones occupied by in-house peptides (yellow) versus peptides from TrEMBL (purple) in Fig. 3 depicts clear separation of in-house peptides from TrEMBL peptides, broadly distributed over the map. This is in line with the fact that only 62 of the 38,642 peptides from TrEMBL were annotated with keywords related to possibly relevant bioactivity (see Supplementary note TrEMBL peptides filtered by keywords), and hence do not overlap much with zones populated by in-house ABPs (Fig. 3). The comparative landscape of in-house peptides and a small fraction of antibiofilm peptides retrieved from the Biofilm-active AMPs database (BaAMPs)^30^ on the right of Fig. 3 shows that the biofilm-targeting peptides cluster near active in-house peptides, clearly delineating zones occupied by antibiofilm peptides. Similarly, zones dominated by in-house peptides with high antibiofilm activity but lower activity against MRSA planktonic cells as defined by the activity threshold chosen in this study can be easily identified (Fig. S2).

**Figure 3.**
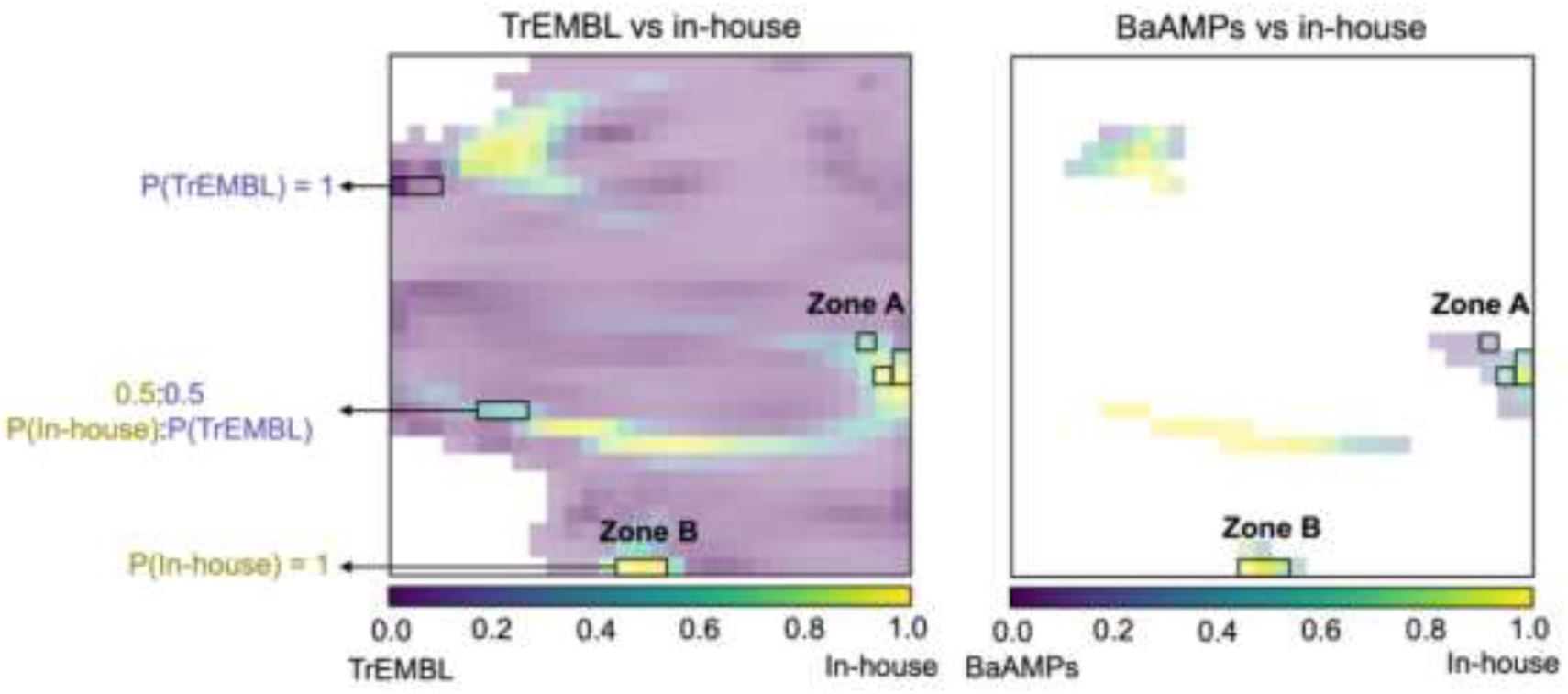
The comparative landscapes reflect the relative node-wise populations of peptides from either of compared databases (TrEMBL/in-house, BaAMPs/in-house). TrEMBL vs. in-house landscape on the left: the purple regions are entirely populated by TrEMBL peptides, while yellow stands for in-house peptides. Right: BaAMPs (purple) vs. in-house (yellow). The nodes of in-between colours (blue, turquoise, green, etc.) are populated by peptides from both databases in varying proportions (e.g., the probability of finding peptides from either TrEMBL or in-house collection is 0.5 in light-blue zones).

### Interpretable generation of peptide analogues from GTM zones

When provided with a latent vector, the trained decoder translates it to an amino acid sequence. To obtain novel latent vectors that would be decoded to more active and diverse analogues of the peptides with already known activity and properties, activity landscapes were used as navigators, enabling the targeted selection of the zones populated with active peptides and subsequent exploration of the peptide space nearby (Fig. 4).

**Figure 4.**
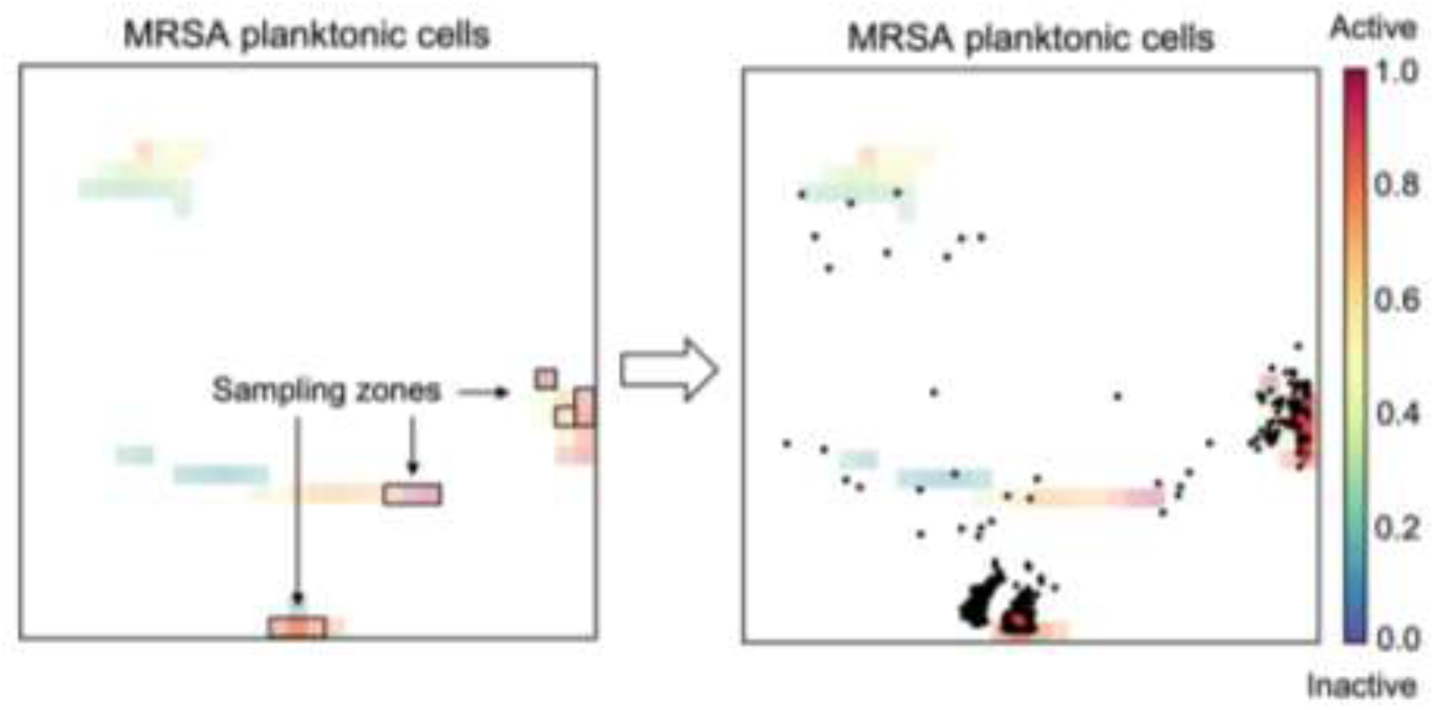
Class landscapes colored by the probability of the active class peptides (inhibiting MRSA planktonic cells) to reside in the corresponding node, see Supplementary Fig. S1 for the anti-MSRA biofilm activity class landscape. The landscape showcases nodes used for sampling in black rectangles, while the barycenter projections of 1,214 generated peptides selected after *in silico* filtering are depicted as black dots. Eight experimentally tested peptides are shown as red points (barycenter projections). Solely the nodes with number of resident peptides higher than one are shown on activity landscapes.

GTM property landscapes are, beyond visualization tools, competent property predictors. The GTM model on latent vectors was compared to Random Forest (RF) and Support Vector Machines (SVM) that were trained on labelled peptide latent vectors to predict activity. All methods performed well in a 5-fold cross-validation repeated five times (Fig. S3 A). Hence, given the good performance of the GTM-based models for activity prediction tasks, generating latent vectors from active zones and then decoding them into amino acid sequences, is anticipated to yield potentially active analogs. To generate novel peptides from a specific area of the map, the coordinates of the GTM nodes were transformed to associated latent vectors that were subsequently used as a mean vector for sampling. A uniform standard deviation of 0.8 was set across all latent dimensions in an attempt to explore the latent space in a proximate vicinity of the node center (mean vector). Hence, 10 nodes from different zones of the map consensually populated by peptides active against both MRSA planktonic cells and biofilms were selected for sampling as shown in black rectangles in Fig. 4. The sampled latent vectors were then decoded to the sequences of amino acids. From each node, 30,000 peptides were sampled, with total number of sampled unique peptides non-identical to the in-house peptides being equal to 298,574.

To narrow the list of peptides for experimental screening, they were encoded to 2-mer vector counts and were filtered based on a 2-mer-based QSAR classifiers applicability domain (AD) and activity prediction confidence criteria (see methods subsection “QSAR models”). In more detail, from the remaining peptides, only those with the probability score to be predicted as active ≥ 0.8 against planktonic cells, as consensually predicted by RF and SVM, were retained. This selection protocol yielded 1,214 peptides predicted to be active against MRSA planktonic cells, among which 1,014 were also predicted to be active against MRSA biofilms.

The aforementioned 1,214 peptides were projected back onto the landscape colored by inhibitory activity against MRSA planktonic cells and appeared to reside primarily in or close to the active zones shown in Fig. 4 right. The tight clusters of designed peptides closely matched the zones from which they were sampled, with eight experimentally tested peptides (test set peptides) shown as red points residing in the centers of the clusters. A minority of developed peptides are more broadly distributed across the map.

### Experimental screening of activity of generated peptides against MRSA

Since the number of peptides to be tested was limited, the eight peptides from the 100 most-active peptides were selected for activity testing, with an emphasis on diversity of amino acid motifs present. This subset of eight were chosen as being the same length as the in-house peptide training set (12-mers) and the control broad spectrum anti-biofilm peptide 1018^22,23^. The eight peptides were tested against both MRSA USA300 planktonic cells and biofilms (Table 1). The peptide IC_50_ values were compared to peptide 1018, which effectively targets a wide range of pathogenic bacteria species, growing as biofilms or to a somewhat lesser extent planktonic bacteria.

**Table 1.**
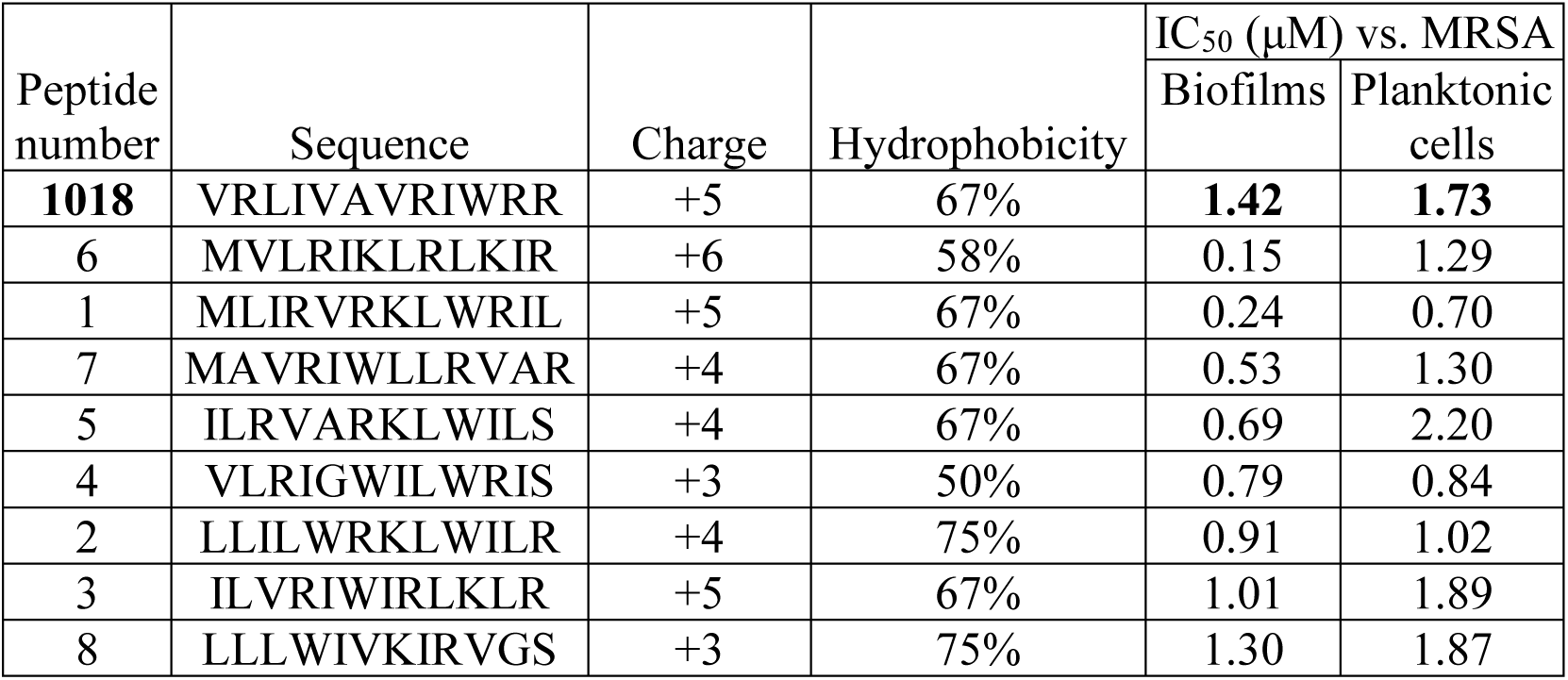
The IC_50_ values calculated based on dose response curves for activity against MRSA biofilms and planktonic cells. The best IC_50_ value out of three repeats for the control peptide is shown. After the control peptide, peptides are ranked according to activity against MRSA biofilms.

Generally speaking, the best antibiofilm peptides had high charge (+4 to +6) and intermediate hydrophobicity (58-67%). Decreasing charge and decreasing or increasing hydrophobicity appeared to reduce activity. Overall, all eight 12-mer peptides (100%) synthesized were active and all were more active against biofilms than positive control peptide 1018. They demonstrated reduced but still good activity against planktonic cells and 5 had better activity than 1018. The most active peptide 6 demonstrated a ten-fold improvement in biofilm activity, when compared to the reference anti-biofilm peptide 1018, while also exhibiting good activity vs. planktonic MRSA. Peptide 1 improved antibiofilm and anti-planktonic activity by 6 and 2.5-fold, indicating broad spectrum improvement. The somewhat lower activities of all peptides against planktonic cells can be attributed to the fact that the probability of the peptides targeting planktonic cells locating in selected nodes was slightly smaller than one as evidenced from the orange node color, whereas the same zones had a probability close to one to be populated by biofilm active peptides (Fig. 4). This was also consistent with the use of anti-biofilm activity as the major activity determinant in the test set of peptides. Overall, the biofilm inhibition hit rate obtained with the introduced pipeline equals or surpasses the 90% hit rate of peptides targeting *E. coli* reported by Vishnepolsky et al.^34^, 83.3% hit rate of membranolytic anti-cancer peptides designed by Grisoni et al.^35^, as well as 10% broad-spectrum AMPs hit rate obtained by Das et al.^27^. Furthermore, the best peptides developed in our study, with IC_50_ activities as low as 0.15 µM, exceeded those obtained in previous ML studies. Further expanding the pool of tested peptides could provide more detailed insights into sequence-activity relationships and potentially lead to the discovery of even more active peptides.

### Minimum biofilm inhibitory concentration, hemolysis and cytotoxicity

The minimum biofilm inhibitory concentration as well as hemolytic activity and cytotoxicity towards peripheral blood mononuclear cells (PBMCs) were measured for peptides 4,6, and 7 obtained at >95% purity (Table 2). All peptides demonstrated high activity as compared to the reference peptide 1018, effectively inhibiting biofilm formation of MRSA USA300. Two peptides (4 and 7) showed selectivity towards *S. aureus*, while peptide 6 was effective against both *S. aureus* and the Gram-negative *P. aeruginosa* PAO1. The biofilm inhibition data for *P. aeruginosa* indicate that distinct peptide features are responsible for the activity in case of peptides 4 and 7, while also referring to some features present in peptide 6 that lead to overlapping activities against both MRSA and *P. aeruginosa* biofilms. Likewise, hemolysis and cytotoxicity values suggest that the peptide features defining toxicity are unique but overlapping with their antibiofilm effects. The peptide toxicity endpoints could potentially be incorporated into the future modelling studies to simultaneously select peptides with low toxicity and high antimicrobial potency.

**Table 2.**
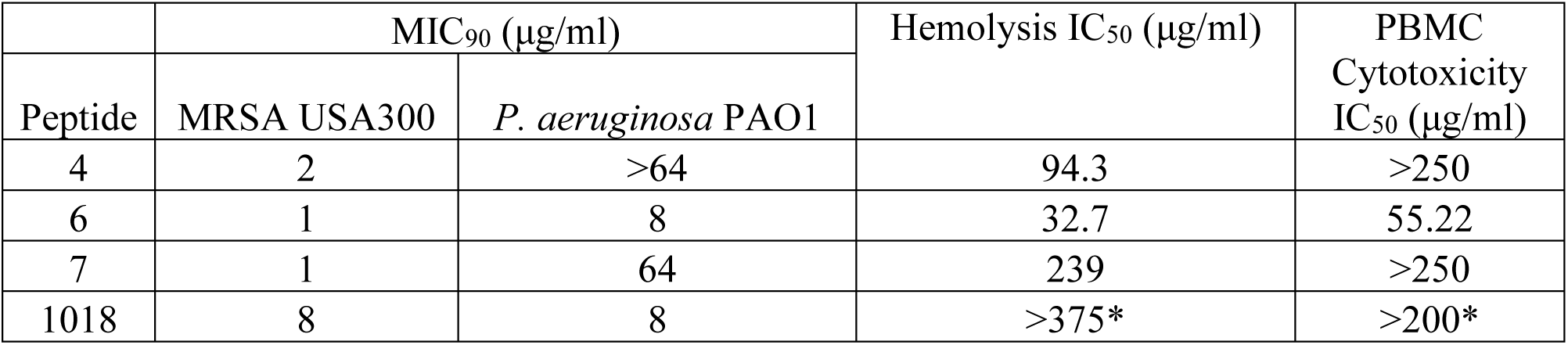
Minimum biofilm inhibitory concentration (MIC_90_, μg/ml) defined as the amount of peptide that inhibited 90% of biofilm growth in a microtitre plate assay, hemolytic activity and cytotoxicity towards PBMCs. Shown is average of three biological replicates from blood cells collected from healthy volunteers. *Hemolytic activity and cytotoxicity towards PBMCs of peptide 1018 were taken from Wieczorek et al.^36^

### Comparison of tested peptides with the in-house sequences

To estimate the similarity of tested peptides to the active in-house set, the analysis of their sequence similarity (based on BLOSUM62 scores) as well as sequence-based calculated properties (Table S2) was conducted. The sequence similarity between tested peptides and in-house peptide collection active against MRSA biofilms and planktonic cells varied from 1 to 12 with mean value being 5, suggesting a moderate level of diversity among the corresponding peptide sets (Fig. S4, Fig. S5).

There were no significant differences in the mean minmax scaled feature values (hydrophobic ratio, Levitt alpha-helix propensity, amino acid transmembrane propensity scale, Eisenberg hydrophobicity, and hydrophobic moment) between the in-house and tested peptide sets as assessed by the pair-wise Welch *t*-test as evidenced by *p*-values > 0.05 (Table S2 and Fig. S6). In contrast, statistically significant differences were found in in-house and tested peptide mean values of the potential estimating energies of amino acid insertion into lipid bilayers (Welch’s *t*-test *p*- value of 0.001). This, along with moderately pronounced amphipathic character of both in-house and tested peptides (overall values of the hydrophobic moment > 0), can be related to the interaction with lipid bilayers and membranes.

## Discussion

Among various methods for the *de novo* design of peptides with antibiofilm and antimicrobial activity, our approach stands out by its ability to render the peptide generation process visually interpretable. This design is aligned with the principles of eXplainable AI (XAI), allowing for human-readable navigation within the latent space. Specifically, the utilization of 2D maps enables manual fine-tuning of the generation process by selecting the most promising regions within the peptide space, as graphically depicted on the maps in Figures 2-4. Moreover, the GTM nodes serve as convenient seeds for sampling, obviating the need to apply additional techniques, such as selecting individual compounds as seeds or performing clustering to identify and use cluster centroids.

Another significant advantage of the GTM-driven approach for *de novo* design is its multi-property prediction capacity. Landscapes related to other relevant properties, such as cytotoxicity, stability, lack of aggregation, etc., can be easily accommodated on the same map. Thus, the current approach can be adapted for sampling, for example, from the regions rich in bioactive yet non-toxic, stable and non-aggregative peptides. The landscapes can be dynamically refined in an active learning manner with newly acquired data, without the need to retrain the GTM, thus addressing applicability domain constraints.

The developed methodology can be further extended for the rational design of cyclic and linear peptides incorporating non-natural amino acids for better affinity, conformational rigidity, and pharmacokinetic properties. Currently, a major limitation is finding a suitable training set of peptides. However, with the advent of high throughput peptide array methods and rapid screening procedures^3,37^ we can cost effectively improve the training set in terms of quality (through standardized activity test methods to avoid discrepancies in assay types used to measure IC_50_) and in terms of quantity of peptides used for training, performing these improvements in an iterative manner. In this regard, it is worth noting that the use of an in-house training set based on antibiofilm activity resulted in peptides with superior activities against MRSA biofilms - the major target for which there are no specifically approved drugs.

## Conclusions

In summary, an end-to-end pipeline for *de novo* design of highly potent antibiofilm peptides was introduced. Based on autoencoder neural network and generative topographic mapping, the introduced approach enables illustrative sampling of novel peptides from the latent space. The developed tool is highly effective, with all tested peptides showing activity against MRSA biofilms as assessed by experimental screening. The experimentally confirmed antibiofilm potency of the generated peptides showcases the utility of the presented pipeline, demonstrating its potential to streamline the decision-making in the drug discovery process.

## Experimental Section

### Data collection and pre-processing

#### Experimental dataset

An in-house set of 680 unique peptides of length 12 was prepared and tested against MRSA following the protocols described below. This data was used as a labelled training set. The distribution of IC_50_ values of the tested peptides was skewed toward ≤ 5µM; hence the threshold was tuned to be stringent dividing the labelled dataset to highly active and moderately active/inactive ones. Namely, peptides with IC_50_ ≤ 5 µM were classified as active, whereas those with IC_50_ > 10 µM were labelled as moderately active/inactive, further referred to as inactive sequences or decoys. To classify peptide sequences with different repeatedly measured IC_50_ values, the mean IC_50_ was computed if standard deviation within the peptide duplicate set was smaller than 10, a condition met by all duplicate sets. Peptides with 5μM < IC_50_ ≤ 10 μM were discarded to provide a more reliable discrimination between highly active and inactive peptides.^29^ These thresholds resulted in active-to-inactive class ratios of 4:1 against planktonic cells and 9:1 against biofilms.

#### Data collection for the autoencoder training

The dataset for the autoencoder training and subsequent use in GTM construction consisted of experimentally tested in-house peptides as well as peptides of lengths 10-14 inclusive collected from TrEMBL, DBAASP, SATPdb, SwissProt, FermFooDb, Hemolytik, NeuroPedia, BaAMPs and APD3 databases (Table S1). The in-house peptides were exclusively composed from proteinogenic amino acids. Peptides with non-natural amino acids from public databases were filtered out. The overall size of the dataset used to train the autoencoder and for GTM projections was hence 45,072. It was divided into the training and validation sets with 80/20 split, comprising 36,057 peptides for training and 9015 for validation.

### Wasserstein AutoEncoder

The implemented architecture was inspired from the results on the peptide generation task with Wasserstein autoencoder^38^ as reported by Das et al.^16^ The WAE encodes each peptide as a distribution in the latent space, with the goal that the mixture of distributions of all peptides from each batch tends to match the prior, i.e., normal distribution. This approach aims to achieve a continuous latent space representation and enhanced disentanglement^39^, facilitating a clearer separation of latent features as opposed to the variational autoencoder (VAE) that may yield more overlapping latent representations. To leverage the ability of the model to learn the plausible peptide grammar, the architecture was trained on a dataset of short sequences of lengths from 10 to 14 amino acids extracted from databases of peptides and proteins both with and without known biological activity across various biological systems. The WAE architecture features a probabilistic (random) encoder equipped with self-attention layers, a gated recurrent unit (GRU) decoder and a linear layer to predict the peptide length from the latent vector. The general scheme of the architecture is presented in Fig. 5. The multi-headed self-attention in the encoder was introduced to learn long-range dependencies across the amino acid sequence. A detailed description of the architecture, underlying layers, and loss functions used in the WAE optimization is provided in the corresponding Supplementary notes.

**Figure 5.**
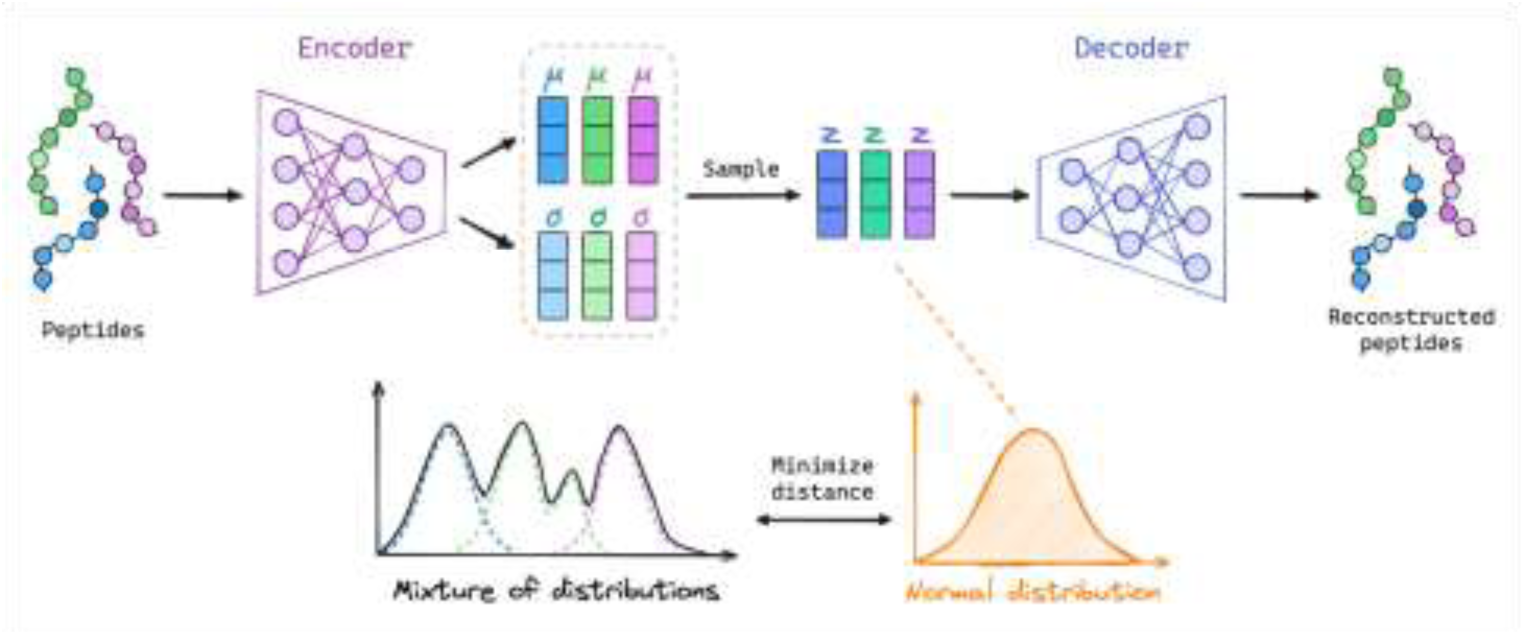
General scheme of WAE architecture applied to peptides.

### GTM

Generative Topographic Mapping (GTM) is a non-linear dimensionality reduction algorithm introduced by Bishop at al.^40^. Its capabilities have been extensively studied in the context of chemical space^41–45^ and biological sequence space exploration^46–48^, chemical database comparison^42^, as a model for property^41,49^ and phenotype^47,50^ prediction, and most importantly, in *de novo* compound design by combining GTM with deep learning models^20^ as well as for the inverse QSAR task^51^. The GTM allows non-linear mapping of the data points in multi-dimensional descriptor space to 2D latent space, represented as a finite squared map of nodes. The mapping is done by inserting a flexible hypersurface called manifold into the multi-dimensional descriptor space to approach the data clouds as closely as possible. The initial step of the GTM, however, consists in the creation of a two-dimensional latent space, i.e., a map, which consists of k nodes with (l_x_, l_y_) coordinates. The mapping of k^th^ node **x_k_** to a corresponding grid point **y_k_** on a flexible manifold is done through a weighted mixture of B non-linear radial basis functions (RBFs), each having the following form:

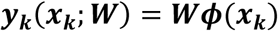

Each of the RBFs is typically a Gaussian whose center is constrained to be taken from the 2D map. Once the mapping from latent space to data space is done through the RBF network (i.e., manifold is initialized), each of the resulting grid points on a manifold becomes the center of the multidimensional Gaussian function describing the distribution of data points in its proximity. The Gaussian function accounting for the spread of data points around the manifold, and reflecting the probability of a data vector ***t***_***n***_ to be assigned to node ***x***_***k***_, has the following form:

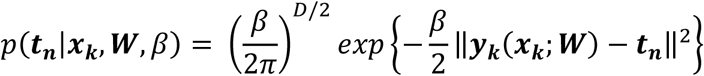

where ***t***_***n***_ is a n^th^ data point vector in a multidimensional space, ***x***_***k***_ is a k^th^ node, ***y***_***k***_(***x***_***k***_; ***W***) is grid point coordinates in multidimensional space, *β*^−1^ is the variance of this Gaussian distribution.

Sampling from this distribution allows for the generation of novel data points that are considered to possess similar characteristics as the neighboring point. The fitting of the manifold to the representative set of data points denoted frame set, i.e., identification of optimal **W** and *β*^−1^, is done with Expectation-Maximization (EM) algorithm by maximizing the log-likelihood (*Llh*) or in simpler terms maximizing the closeness of the manifold to the data points. Namely, the *Llh* is defined as an estimate of how well the manifold fits each point of the frame set ***t***_***n***_ to every node

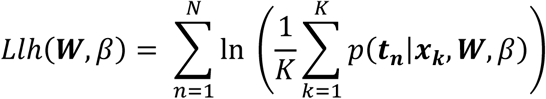

Once the manifold is optimized, the (new) data points can be projected onto it, resulting in a vector of responsibilities for each data point which represent probabilities of each data point to be projected into each node of the map:

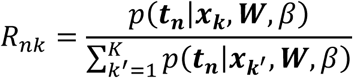

Upon node-wise summation of responsibilities across the map, fuzzy density landscapes can be created that are “colored” according to the extent of the node population. In a similar way, the map can be colored according to the properties of the data points (peptide sequences) or the probability to find sequences of a certain class residing in a given node, yielding property or class landscapes respectively. The transparency added to the class or property landscapes reflects the population density of the nodes.

The landscapes facilitate the analysis of the distribution of sequences across the map and the identification of patterns related to the spread of sequences with specific features. This includes identifying areas abundant in peptides that share similar bioactivity profile, zones predominantly containing hydrophobic peptides, or the ones populated by peptides with certain amino acid motifs, to name a few. Such analysis aids the selection of optimal zones for generation of focused peptide libraries.

In this work, the GTM manifold was fitted to 10,000 randomly chosen peptides from the collected dataset encoded to latent vectors with WAE. To have a complete representation of the latent space of the collected peptides, all the 45,072 sequences were projected onto the manifold, generating a density landscape. To generate activity landscapes, the population of nodes was complemented by the underlying class labels of the in-house peptides. Only in-house activity data was used to avoid the bias induced by discrepancies in assay types used to measure IC_50_ when working with labelled data from multiple sources.^52^

#### GTM-based predictive models

Apart from pure visualization, the landscapes also served as predictive models. The new sequences, encoded using the same descriptors as the initial dataset, can be projected onto the map, with prediction value being deduced from mean property values or probabilities to belong to a class of nodes it was projected to. The predictive propensity of the underlying landscape-based models is directly related to the quality of the manifold. Namely, apart **W** and *β*^−1^, the manifold is defined by its hyperparameters: number of map nodes, number of RBFs, common width of the RBFs, and a regularization coefficient. As opposed to unsupervised manifold fitting, the selection of optimal hyperparameters as well as descriptor sets for data representation is achieved by maximizing an objective function, herein - predictive propensity of the given landscape in classification or regression tasks. To find the optimal hyperparameters, the Genetic Algorithm (GA) was used. The GA is stochastic evolutionary algorithm allowing browsing through the space of GTM hyperparameters and subsequent choice of Pareto-optimal ones by maximizing Balanced Accuracy (BA) in predicting peptide activity label. In more detail, herein, the GA optimized hyperparameters of the GTM using unprocessed WAE latent vectors as a fixed set of descriptors. To evaluate the GTM’s predictive accuracy for class landscapes, a balanced accuracy was computed from 5-fold cross-validation (5-CV) repeated five times with reshuffled sets. In each iteration, 80% of the dataset was used as a training set to color the map, while the remaining 20% served as a test set for projection of the corresponding data points onto the landscape, and performance evaluation.

### QSAR models

The Random Forest (RF) and Support Vector Machine (SVM) classification models implemented in the scikit-learn library (v. 1.1.1) were built on peptide sequences represented as vectors of 2-mer counts with experimentally measured IC_50_ values. The 2-mer counts vectors were subjected to feature scaling implemented with StandardScaler of scikit-learn library (v. 1.1.1). The hyperparameters were tuned by grid search optimization algorithm. To assess the predictive performance of the developed QSAR models, the five-fold stratified cross validation was performed. The performance of the SVM and RF models in predicting activity against MRSA planktonic cells resulted in BA values of 0.721 and 0.742 respectively, whereas for MRSA biofilms, the SVM and RF models achieved BAs of 0.736 and 0.759, respectively (Fig. S3 B). The formula of BA calculation used to evaluate the performance of the models is reported in supplementary note on statistical metrics for evaluation of QSAR models. The rationale behind creating classification models instead of regression ones was due to the skewed distribution of IC_50_ values (see Data collection and pre-processing section for details). The activity labels of 298,574 generated peptides were predicted with created QSAR models. To ensure the reliability of activity predictions for generated peptides, only those peptides that fell within or near the bounds of the models’ Bounding Box (BB)-based AD^53^ were considered. The AD was defined as a hyper- rectangle bounded by minimum and maximum values of each of the k-mers’ counts descriptors from the training set of the QSAR model. Peptides whose 2-mer vectors extended outside the AD bounds in more than two dimensions were excluded. This process resulted in two peptide sets: 4,563 peptides that fulfilled AD criteria defined by QSAR models predicting activity against MRSA biofilms, and 4,578 within AD defined by QSAR models predicting activity against MRSA planktonic cells. All 4,563 peptides from the first set were also included in the latter set of 4,578 peptides. Further filtering was applied to the latter set of 4, 578 peptides. Namely, they were ranked based on classifier’s prediction confidence. Those peptides that were predicted to be active against planktonic cells by RF and SVM models with probability score higher than 0.8 were kept. Overall, it resulted in 1,214 peptides, from which 1,014 were also predicted to be active against biofilms with probability score higher than 0.8 by RF and SVM models.

### Peptide synthesis and stock solution preparation

Peptide arrays on cellulose membranes were made by Kinexus Bioinformatics Corporation (Vancouver, BC, Canada) and were obtained as cleaved free peptide associated with the punched-out spot of the cellulose membrane. Stock solutions of the peptide array samples were prepared by dissolving the free peptide in 200 µl of endotoxin free sterile water, incubating at room temperature for ∼2 hr, and then transferring the resulting peptide solution to a sterile microfuge tube for subsequent antimicrobial activity assays. The concentration of the stock peptide array samples was determined by measuring the absorbance at 280 nm of the solution (in duplicate) on a BioTek Epoch Microplate Spectrophotometer using the Take3^TM^ Microvolume Plate. A theoretical extinction coefficient of each peptide was determined based on the number of Trp and Tyr residues in the peptide sequence according to the formula: (#Trp) * 5500 M^-1^cm^-1^ + (#Tyr) * 1490 M^-1^cm^-1^, and the concentration of the peptide in solution was calculated according to the Beer-Lambert law. Any Cys residues within the peptide samples were assumed not to be present in a disulfide bond, so their contribution to the extinction coefficient at 280 nm was ignored. Stock concentrations of peptides lacking a Trp and Tyr residue were assigned the average concentration of all the samples for which a concentration could be determined within the same peptide array, which was 392.8 µM.

Peptides synthesized to >95% purity (peptides 4, 6 and 7) were obtained from Peptide 2.0 (Chantilly, VA). Purified peptides were amidated at their C-terminus and were obtained as acetate salts. Peptides were purified by reverse phase HPLC and peptide identity was confirmed by mass spectrometry.

### *In vitro* screening of novel peptides

To validate the peptides generated by WAE and selected with *in silico* methods, a set of eight peptides were selected and SPOT-synthesized on cellulose arrays. The antibiofilm activity of the set of eight peptides was assessed against MRSA USA300 planktonic cells and biofilms using the crystal violet assay.

### Antibiofilm activity screen

The antimicrobial activity and biofilm inhibition activity of the peptide array samples was assessed against methicillin resistant *Staphylococcus aureus* (MRSA) as described previously^54^. Briefly, two-fold serial dilutions of the peptide array samples (10 µl final volume) were prepared in 96- well plates and then 90 µl of an overnight culture of MRSA diluted to an OD_600_ = 0.01 in 10% tryptic soy broth (TSB) supplemented with 0.1% glucose was added to each well. The microtitre plates were then incubated overnight at 37°C. The following day, bacterial growth was quantified by measuring the optical density at 600 nm (OD_600_) of each sample well, while the amount of biofilm was quantified by rinsing the wells of the plate with distilled water and staining the adhered biomass with 0.1% crystal violet solution. The amount of adhered biomass was quantified by recording the absorbance at 595 nm (A_595_). All sample measurements were performed on a BioTek Epoch Microplate Spectrophotometer. Dose response curves were drawn for each peptide based on the bacterial growth (OD_600_) or biofilm inhibition (A_595_), normalizing the percent of peptide induced inhibition relative to untreated bacteria (100% growth) or sterile media (0% growth) growing within the same microtitre plate. The concentration of peptide required to inhibit 50% of bacterial or biofilm growth (IC_50_) was calculated by non-linear regression fit of the dose-response curves using the N-Parameter Logistic Regression (nplr) package^55^ in R^56^.

### Biofilm Inhibition

For the biofilm inhibition experiments of >95% pure peptide samples against MRSA USA300 and

*P. aeruginosa* PAO1, the experimental setup was essentially the same as the spot-synthesis peptide array samples except that PAO1 biofilms were grown in BM2 minimal media (62 mM potassium phosphate buffer, pH 7.0, 7 mM (NH_4_)_2_SO_4_, 0.5 mM MgSO_4_, and 0.4% glucose). Data shown correspond to the peptide concentration that inhibited >90% of biofilm growth based on crystal violet staining of the adhered biofilm biomass within sample well. All biofilm inhibition experiments were performed in triplicate.

### Cytotoxicity and Hemolysis

Cells used in the peptide toxicity assays were isolated from venous blood donated by healthy donors with informed consent in accordance with the University of British Columbia ethics certification and guidelines (approval number H21-00727) which specified the methods used.

Peptide cytotoxicity was assessed against peripheral blood mononuclear cells (PBMCs) using the Cytotoxicity Detection Kit (Roche Diagnostics) which measures the enzyme activity of lactate dehydrogenase (LDH) released from damaged cells^57^. PBMCs were isolated from blood by density centrifugation using Lymphoprep Density Gradient Medium (STEMCELL Technologies) and then resuspended in RPMI media supplemented with 10% fetal bovine serum. Cells and peptide were incubated overnight at 5% CO_2_ and 37 °C and the following day, supernatants were collected and subjected to the LDH assay. Supernatants of cells treated with vehicle (water) or lysed with 2% Triton-X100 (added 1-hr prior to collection of sample supernatant) were used as the negative (0% cytotoxicity) and positive (100% cytotoxicity) controls, respectively.

Peptide induced hemolysis was performed as described previously^58^. Briefly, red blood cells (RBCs) were rinsed three times in PBS and then the hematocrit (100% RBC) was used to prepare an RBC suspension of which 90 µl was seeded into the wells of a 96-well microtitre plate. Peptide treatments or vehicle controls (10 µl) were added to the RBCs and then the plates were incubated for four hours at 5% CO_2_ and 37 °C. The final RBC concentration in each well was 2.5%. Following incubation, the samples were centrifuged (1400 rpm) and the absorbance of the collected supernatants was recorded at 490nm on a BioTek Epoch Microplate Reader. Supernatants of RBCs treated with vehicle (water) or lysed with 2% Triton-X100 (added at the same time as the peptide treatments) were used as the negative (0% toxicity) and positive (100% toxicity) controls, respectively.

## Author contributions

K.P. and T.A. designed the pipeline and implemented the WAE. K.P. performed the computational experiments. K.P., A.O. and T.A. analyzed the results. E.F.H., N.A. and R.E.W.H. conducted the experiments. A.V. and A.C. co-designed and supervised the study. All authors reviewed the manuscript.

## Abbreviations

5-CV: Five-Fold Cross-Validation
ABP: Antibiofilm Peptide
AD: Applicability Domain
AMP: Antimicrobial Peptide
AMR: Antimicrobial Resistance
BA: Balanced Accuracy
BLOSUM: BLOcks SUbstitution Matrix
EM: Expectation Maximization
FDA: Food and Drug Administration
GA: Genetic Algorithm
GRU: Gated Recurrent Unit
GTM: Generative Topographic Mapping
HPLC: High-Performance Liquid Chromatography
IC50: Half-Maximal Inhibitory Concentration
LDH: Lactate Dehydrogenase
LSTM: Long Short-Term Memory
MDR: Multidrug Resistance
ML: Machine Learning
MRSA: Methicillin-Resistant *Staphylococcus Aureus*
PBMC: Peripheral Blood Mononuclear Cell
QSAR: Quantitative Structure-Activity Relationships
RBF: Radial Basis Function
RF: Random Forest
RNN: Recurrent Neural Network
SVM: Support Vector Machines
VAE: Variational AutoEncoder
WAE: Wasserstein AutoEncoder
XAI: eXplainable Artificial Intelligence.

## ASSOCIATED CONTENT

### Data availability statement

The datasets from the open-source databases collected in the current study are available from the following repository: https://huggingface.co/datasets/karinapikalyova/peptides/tree/main. The experimentally annotated data is available from the corresponding authors A.V. and A.C. on reasonable request.

### Code availability statement

The code for the WAE implemented in this study is available in the GitHub repository (https://github.com/Laboratoire-de-Chemoinformatique/GTM_WAE).

### Supporting Information

Additional activity and comparative landscapes, benchmarking results for models built on both latent vectors and 2-mers counts descriptors, the distributions of BLOSUM62 scores between the tested peptides and in-house peptides, comparison of main peptide features, training set composition, the mean and standard deviation values of unscaled descriptors as well as the associated p-values computed with pairwise Welch’s t-test based on min-max scaled features’ values, biological activity details based on keywords for peptides from TrEMBL, in-depth explanations of the WAE architecture.

## Additional figures

**Figure S1.**
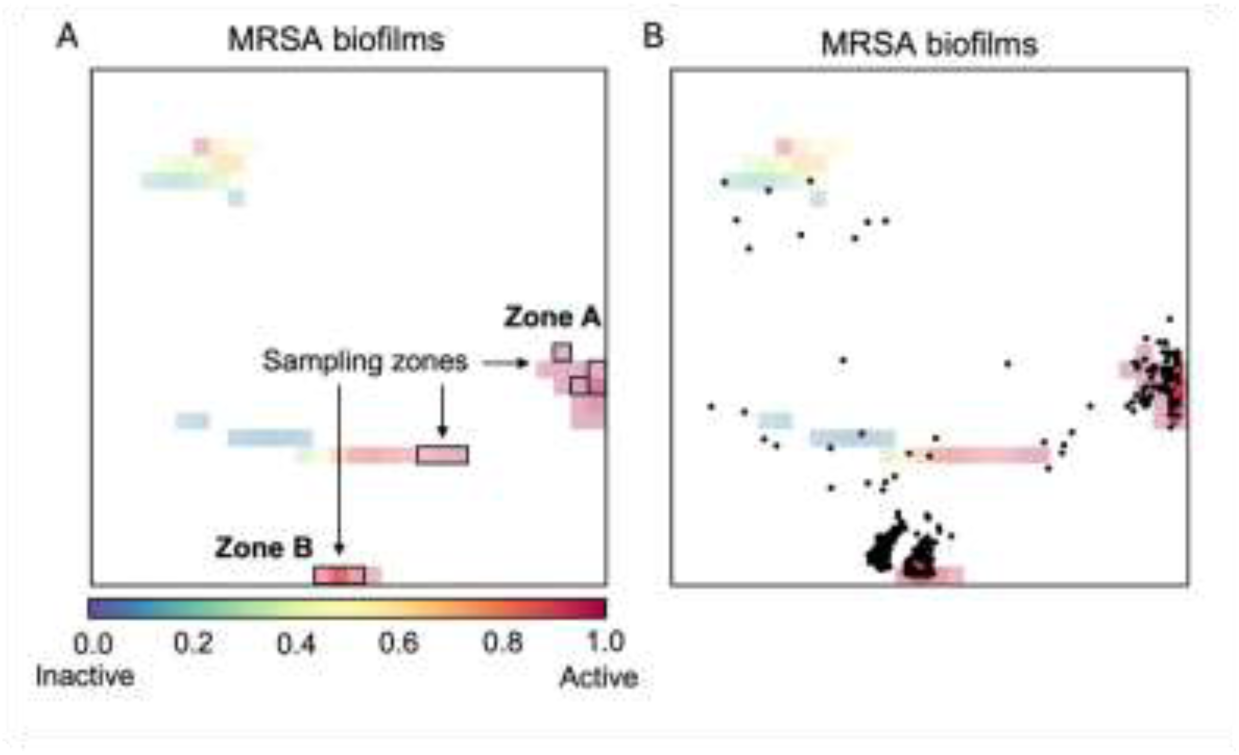
A) Activity class landscape coloured by the probability that active class peptides (inhibiting MRSA biofilms) reside in the corresponding node. The landscape showcases nodes used for sampling in black rectangles. The nodes densely populated by active peptides are denoted zones A and B. B) Anti-MRSA biofilm activity landscape with the projections of 1,214 generated peptides, selected after *in silico* filtering, depicted as black dots. Eight experimentally tested peptides are projected as red points.

**Figure S2.**
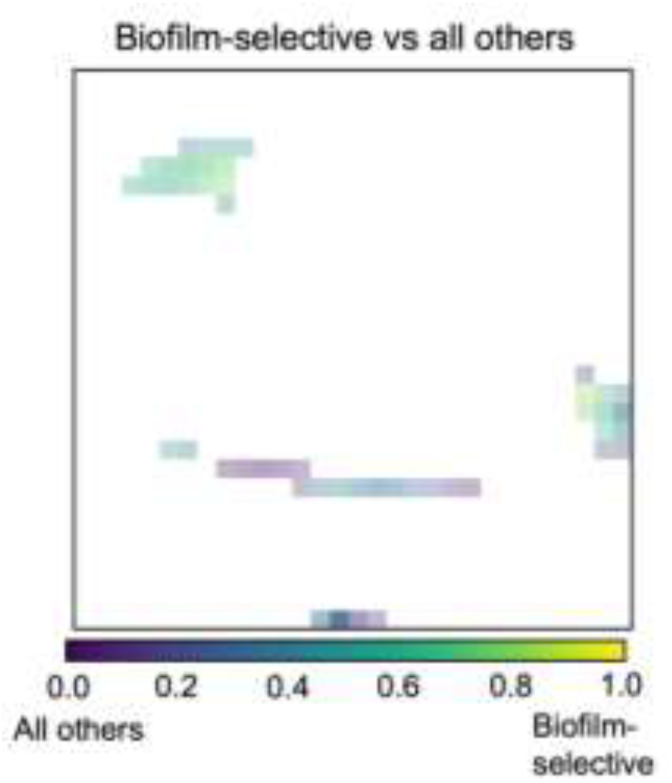
The comparative landscapes reflect the relative node-wise populations of peptides from either of the compared classes. Herein, the class « biofilm-selective » was assigned to peptides defined to be active only against biofilms according to the activity threshold: peptides with IC50 ≤ 5 µM were classified as active, whereas those with IC50 > 10 µM were labelled as inactive. The biofilm-selective peptides are shown in yellow, while all other peptides (recognized as inactive against biofilms or active against both planktonic cells and biofilms) are shown in purple. The nodes of in-between colours (blue, turquoise, green, etc.) are populated by peptides of both classes in varying proportions.

**Figure S3.**
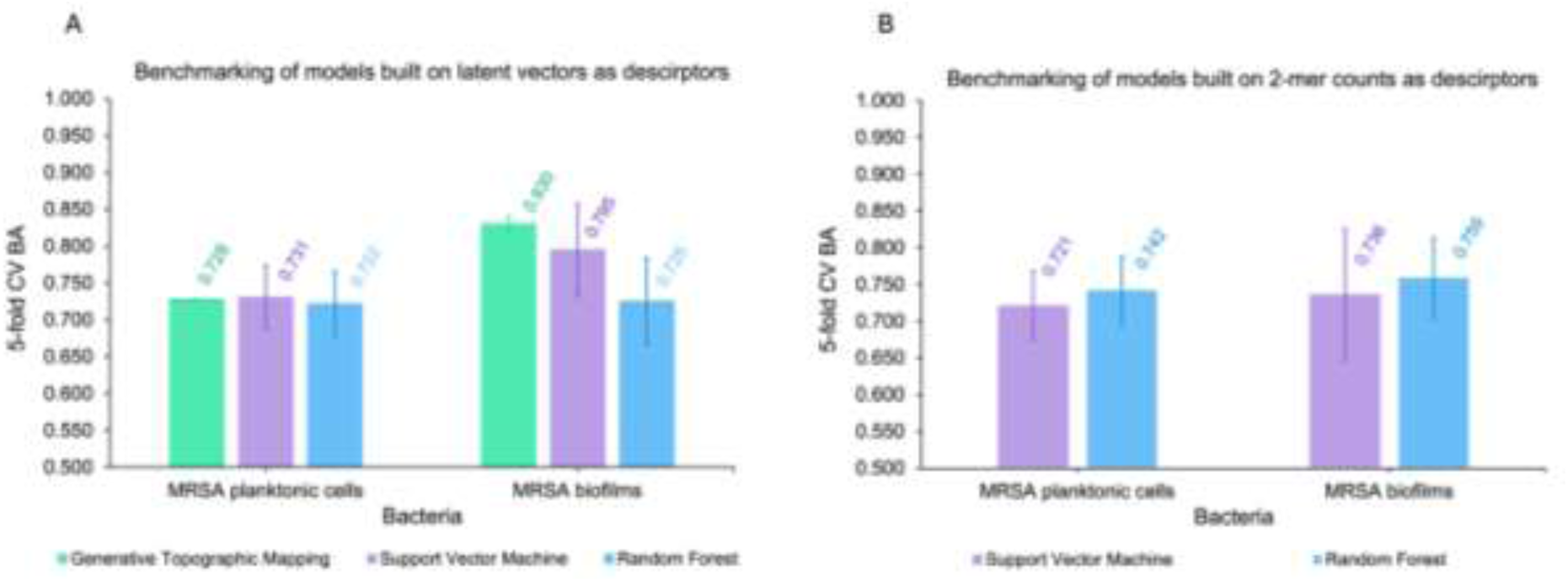
A) The 5-fold cross-validated balanced accuracy values after five repeats for predictions of activity against planktonic cells and biofilms respectively by Random Forest (RF), Support Vector Machine (SVM) and GTM with latent vectors as descriptors. B) The 5-fold cross-validated balanced accuracy values after five repeats for predictions of activity against planktonic cells and biofilms respectively by Random Forest (RF) and Support Vector Machine (SVM) with 2-mer counts as descriptors.

**Figure S4.**
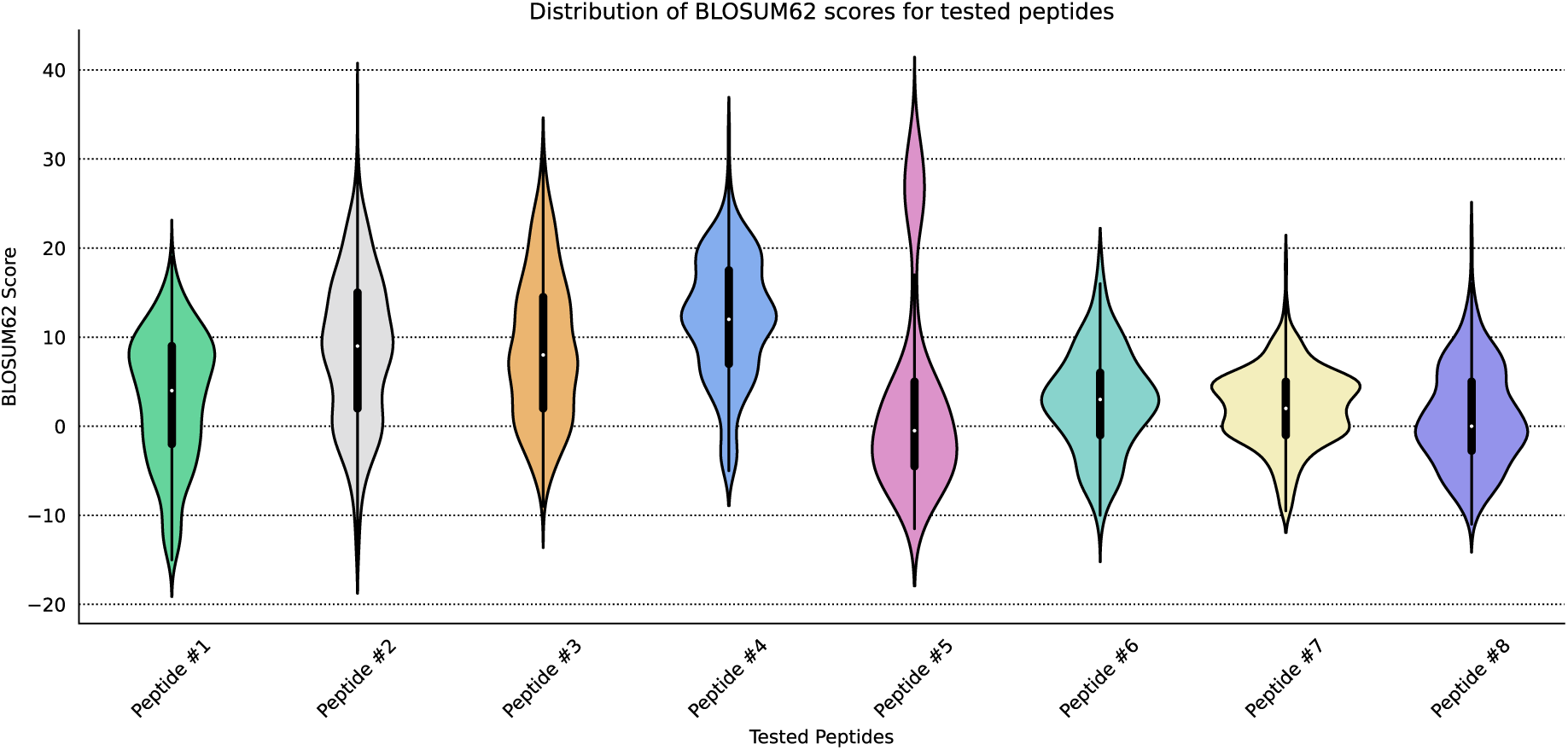
The distributions of BLOSUM62 scores between the tested peptides and in-house peptides active against MRSA biofilms.

**Figure S5.**
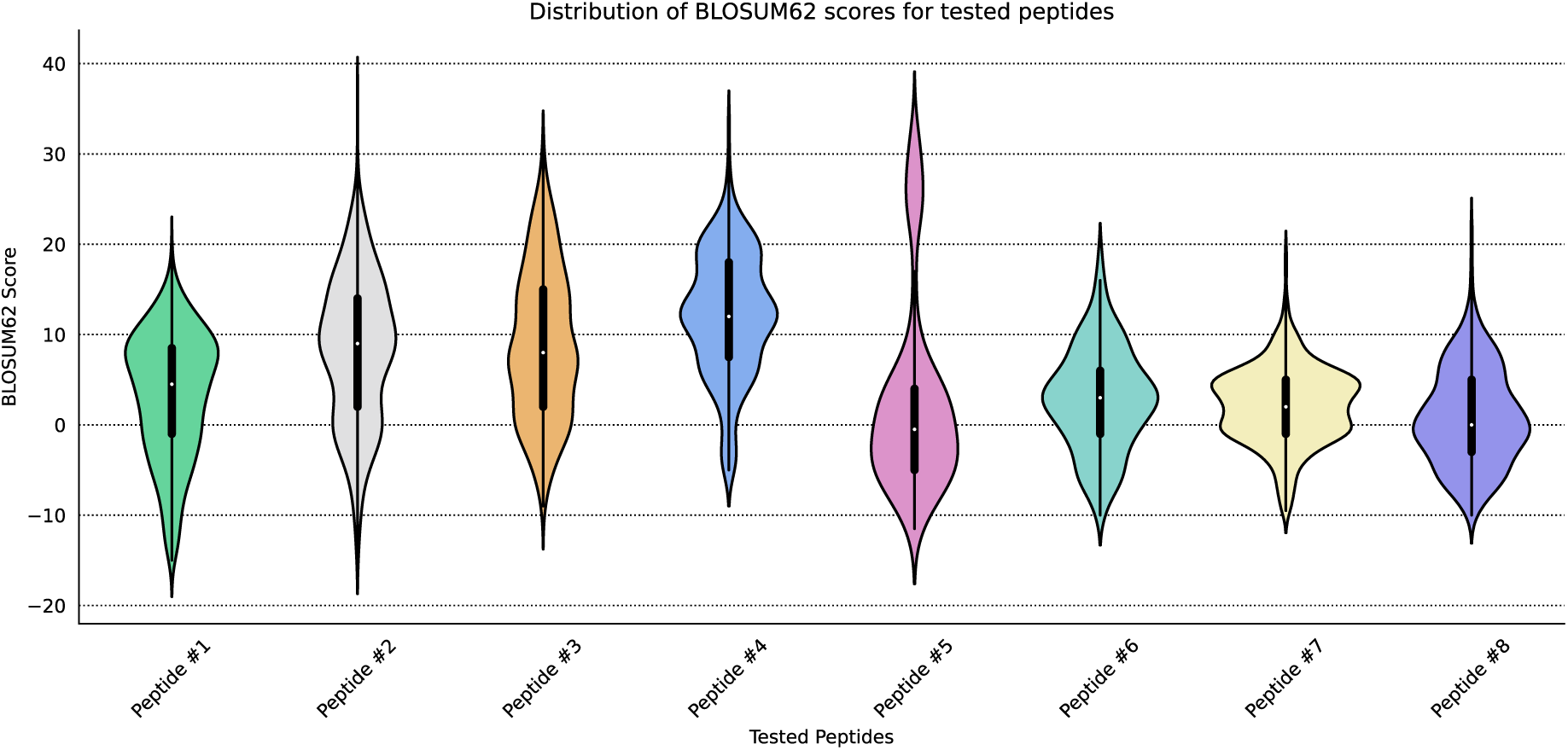
The distributions of BLOSUM62 scores between the tested peptides and in-house peptides active against MRSA planktonic cells.

**Figure S6.**
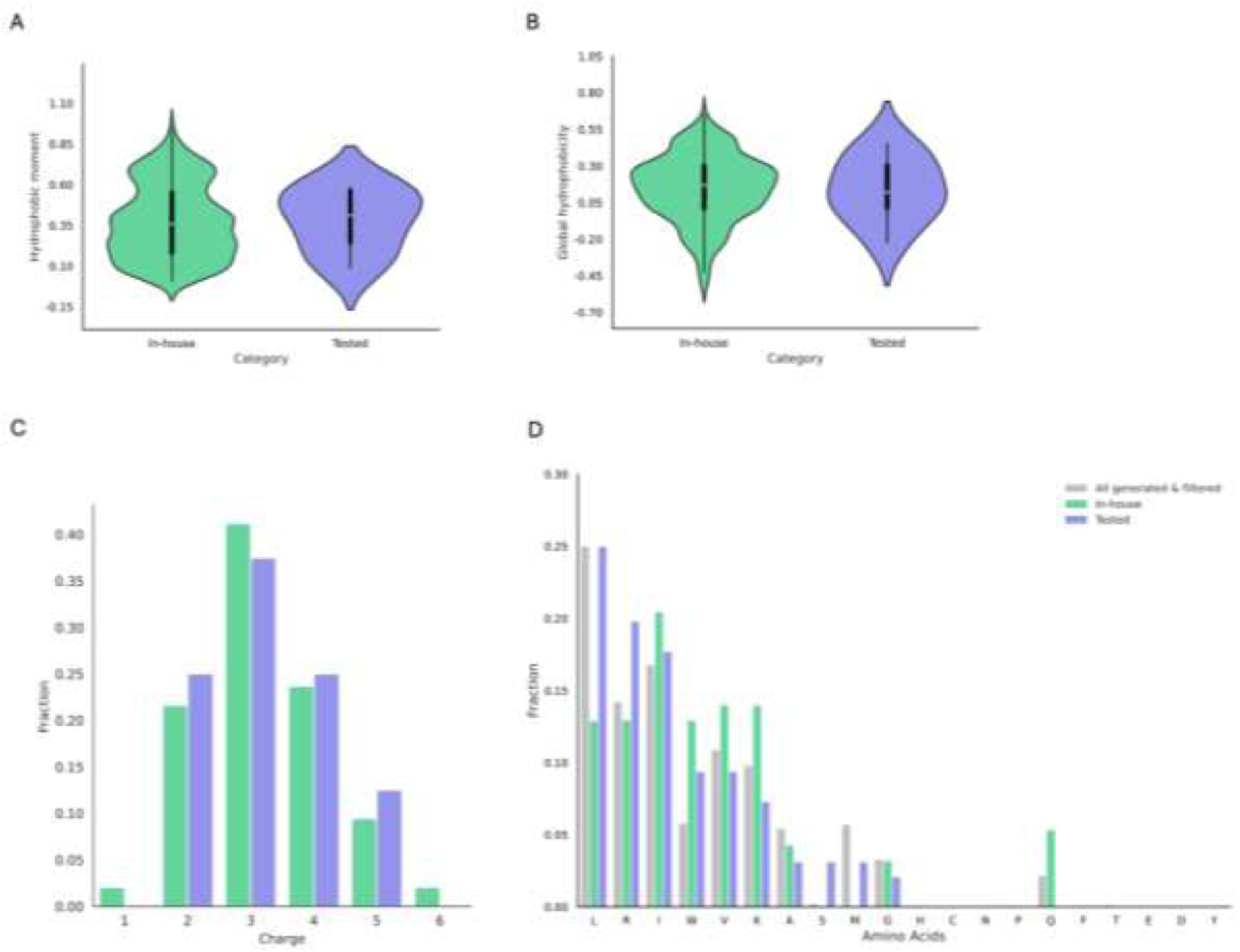
A) Eisenberg hydrophobic moment. B) Eisenberg hydrophobicity. C) Total charge. D) Distribution of amino acids.

## Additional tables

**Table S1.** The composition of the training set.

| Databases | Number of peptides |
| --- | --- |
| TrEMBL | 38642 |
| DBAASP | 3506 |
| SATPdb | 1236 |
| SwissProt | 691 |
| In-house | 680 |
| FermFooDb | 140 |
| Hemolytik | 73 |
| NeuroPedia | 41 |
| BaAMPs | 40 |
| APD3 | 23 |

**Table S2.** Mean and standard deviation values of unscaled descriptors. The associated p-values were computed with pairwise Welch’s t-test based on min-max scaled features’ values. Ez stands for potential evaluating energies of insertion of AA side chains into lipid bilayers, TM is AA transmembrane propensity scale, hydrophobic ratio stands for relative frequency of A, C, F, I, L, M, V, W and Y AAs. All properties were computed with modlAMP package^1^.

| Feature | In-house | Tested | *p*-value |
| --- | --- | --- | --- |
| Eisenberg hydrophobicity | 0.15 ± 0.22 | 0.14 ± 0.22 | 0.914 |
| Eisenberg hydrophobic moment | 0.38 ± 0.23 | 0.39 ± 0.19 | 0.919 |
| Charge | 3.23 ± 1.02 | 3.25 ± 1.04 | 0.957 |
| Hydrophobic ratio | 0.65 ± 0.07 | 0.68 ± 0.05 | 0.141 |
| Levitt alpha helix propensity | 1.06 ± 0.04 | 1.08 ± 0.06 | 0.352 |
| Ez | 18.93 ± 0.88 | 20.27 ± 0.74 | 0.001 |
| TM | 0.14 ± 0.31 | 0.36 ± 0.29 | 0.064 |

## Supplementary Notes

### TrEMBL peptides filtered by keywords

From 38642 peptides of lengths 10-14 collected from TrEMBL, only 62 had one or several of the following keywords: antibiotic, toxin, secreted, antimicrobial, disulfide bond, lipid-binding, membrane, cytolysis, cell wall biogenesis/degradation, amphibian defense peptide, defensin, antiviral protein, fungicide, allergen, hypotensive agent, immunity, innate immunity, inflammatory response, lantibiotic, toxin, neuropeptide, protease inhibitor, cell wall, antiviral defense, antioxidant, bacteriocin, cardiotoxin, cell adhesion impairing toxin, calcium channel impairing toxin, calcium-activated potassium channel impairing toxin, chloride channel impairing toxin, complement system impairing toxin, dermonecrotic toxin, enterotoxin, fibrinogenolytic toxin, fibrinolytic toxin, g-protein coupled acetylcholine receptor impairing toxin, g-protein coupled receptor impairing toxin, acetylcholine receptor inhibiting toxin, blood coagulation cascade activating toxin, blood coagulation cascade inhibiting toxin, bradykinin receptor impairing toxin, platelet aggregation activating toxin, hemorrhagic toxin, hemostasis impairing toxin, ion channel impairing toxin, myotoxin, neurotoxin, pharmaceutical, platelet aggregation inhibiting toxin, plant defense, postsynaptic neurotoxin, potassium channel impairing toxin, presynaptic neurotoxin, proton-gated sodium channel impairing toxin, ryanodine-sensitive calcium-release channel impairing toxin, viral exotoxin, voltage-gated calcium channel impairing toxin, voltage-gated chloride channel impairing toxin, voltage-gated potassium channel impairing toxin, voltage-gated sodium channel impairing toxin, acetylcholine receptor inhibiting toxin.

### Encoding

The input peptides, i.e., sequences encoded with one letter amino acid (AA) code, are first tokenized, padded according to the longest peptide in terms of tokens and then each AA (token) is encoded into embedding vector of dimension *E* (*E =* 256). To conserve the information on AA order within the peptide, the positional encoding matrix of similar dimensionality is added to the embedding matrix. As a result, each peptide is represented as a matrix ***X***: (*x̅*_1_, *x̅*_2_, …, *x̅*_*i*_) where *x̅*_*i*_ stands for the embedding vector of the i^th^ token of the peptide. The matrices ***X*** denoting each peptide serve as input to the encoder. The encoder consists of six stacked self-attention blocks (SelfAttnBlock). All these steps are illustrated in Fig. S7.

**Figure S7.**
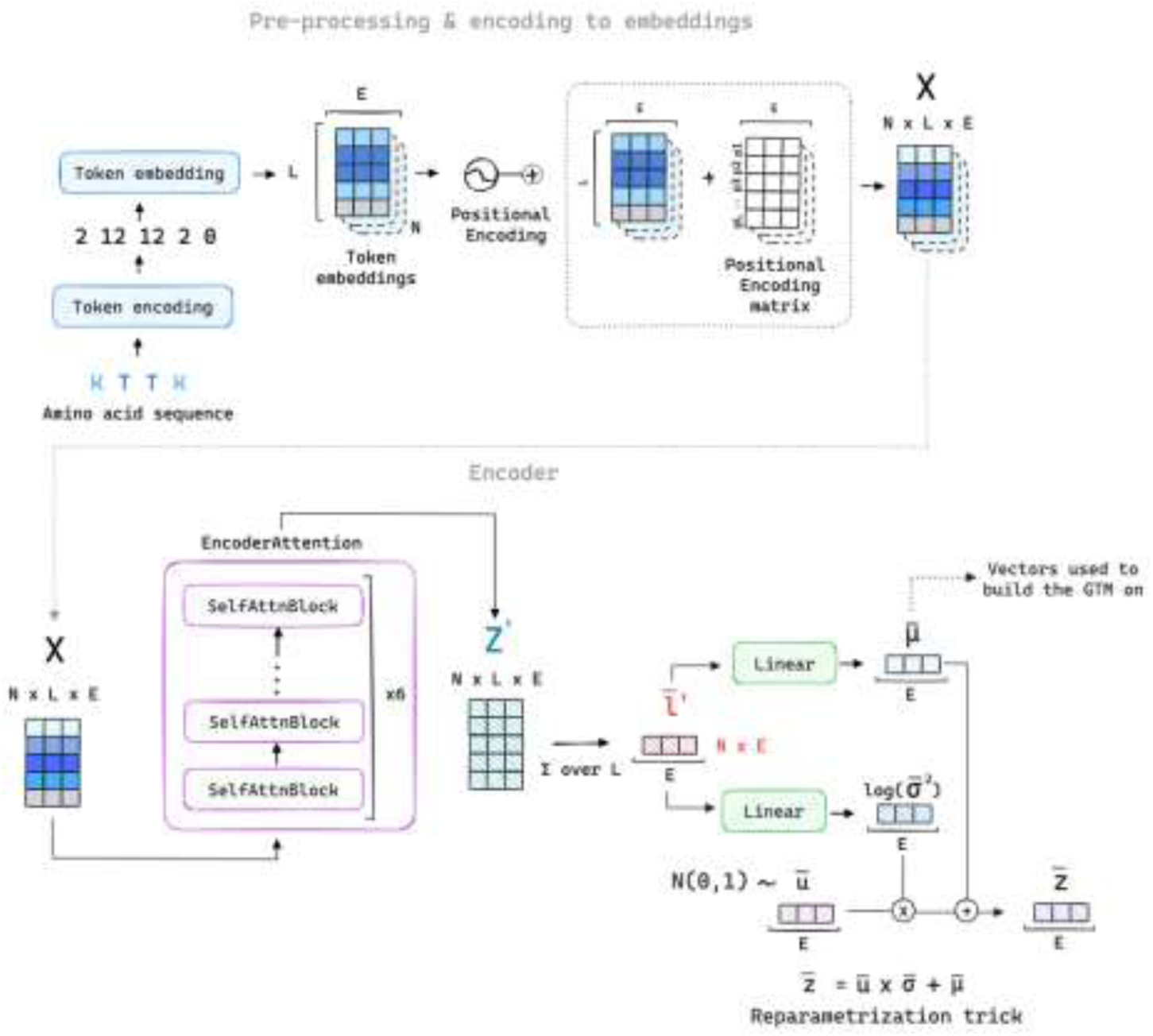
Scheme of pre-processing, encoding to embeddings and the WAE encoding process.

Each self-attention block follows the original transformer encoder layer structure as introduced by Vaswani et al.^2^, with two sub-layers: multi-head self-attention (MHA) and feed forward layers implemented in the PyTorch package. Akin to the original transformer encoder layer, around each sublayer the residual connections are added, with only difference being the layer normalization block being positioned before the MHA layer and after the combination of MHA and dropout layers. The MHA is composed from eight attention heads. The self-attention block and MHA layer are schematically illustrated in Fig. S8.

**Figure S8.**
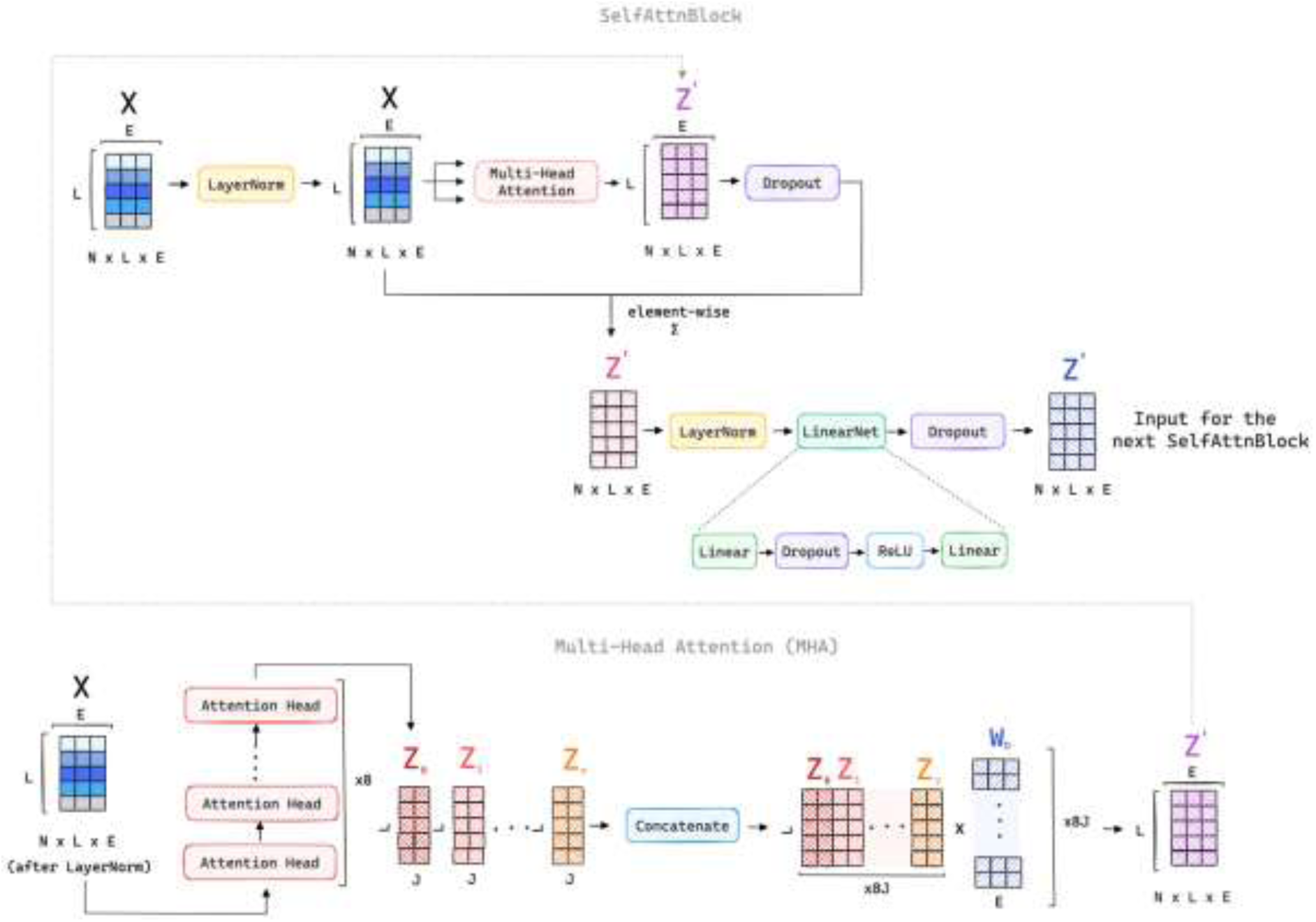
Scheme of the self-attention block and MHA layer implemented in WAE encoder.

Each attention head is used to train weights that would define the degree of relatedness (degree of “attention paying”) of a current AA to all AAs in a peptide. To obtain the attention weights, first, the attention scores *s*_*ij*_ defined as a scalar product of query encoding of the current AA vector - *q̅*_*i*_ with key encodings *k̅*_*j*_ of all AAs’ vectors from the peptide are computed:

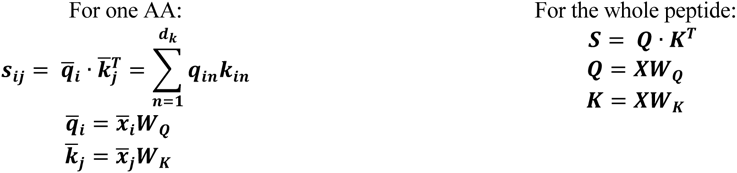

where *q̅*_*i*_ and *k̅* vectors are query and key encodings of *x̅* and *x̅* respectively, ***W_Q_*** and ***W_K_*** matrices are learnable parameters of the self-attention layer; ***Q*** and ***K*** are matrices with query and key vectors for each AA in a peptide. The attention scores allow the comparison of the current AA with all AAs in the peptide. The scores are scaled by √*d*_*k*_ (the square root of dimension of *k̅*_*j*_ vector) to achieve unit variance, and softmax is applied to obtain attention weights *a*_*ij*_ :

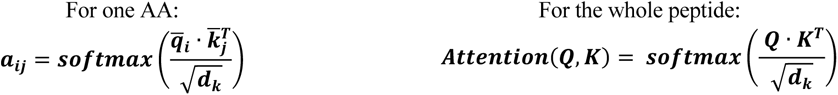

To produce the final encoding *z̅*_*i*_, the attention-weighted sum of value vectors *v̅*_*j*_ of all AAs within the peptide is computed, forming a convex combination:

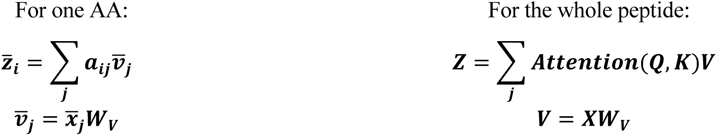

where *v̅*_*j*_vector is a value encoding of *x̅*_*j*_, ***W***_***v***_ matrix is a learnable parameter of the self-attention layer; ***v*** is a value matrix with value vectors for each AA in a peptide. The scheme of the operations done in attention head module is given in Fig. S9.

**Figure S9.**
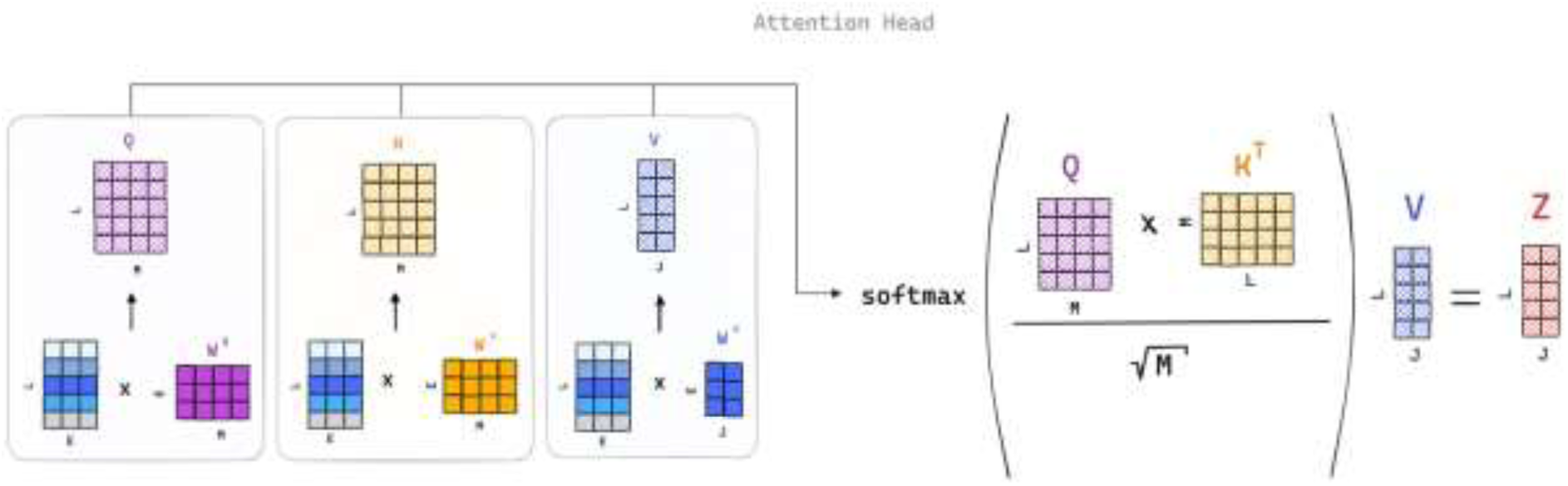
Operations happening in an attention head.

The *z̅*_*i*_vectors after each attention head are then concatenated and multiplied by the **Wo** weight matrix (Fig. S8) to produce the final output vector *z̅_i_*^′^, in the same manner the ***Z*** matrices after each head are concatenated and multiplied by the **Wo** weight matrix. This output is then element-wise summed with the matrix ***X***, and the resulting matrix is passed through the feedforward layer. The resulting ***Z***′ matrix composed from *z̅*^′^ vectors, becomes the input of the next of six self-attention blocks. The final matrix ***Z***′ after all attention blocks (Fig. S7) is summed across sequence length (*L*) direction to get a vector *l̅*′. The latter serves as input to feed forward layer to get mean *μ̅* and variance *σ̅* of the Wasserstein posterior distribution, hence representing each peptide as a distribution in the 256-dimensional space. The reparameterization trick is then done to yield the final *z̅* used for training. The centre of distribution vector *μ̅* is used as descriptor vector for each peptide for the GTM construction.

### Decoding

The obtained *z̅* vector, along with the AA embedding matrix for each peptide are passed to the decoder. Namely, to prepare the input matrix for the GRU, the *z̅* is appended atop the embedding matrix. It is followed by expansion of the *z̅* vector to form a matrix and its concatenation with the modified embedding matrix along the embedding dimension *E.* The formed input feature matrix is hence of dimensionality (*L+1*, *2*E*). The output matrix of the unidirectional GRU layer is then trimmed to remove the last row due its non-viability for token prediction. The trimmed matrix is then passed through a linear layer which outputs the unnormalized probabilities (logits) of each vocabulary token to be found in a given position of the sequence (Fig. S10).

**Figure S10.**
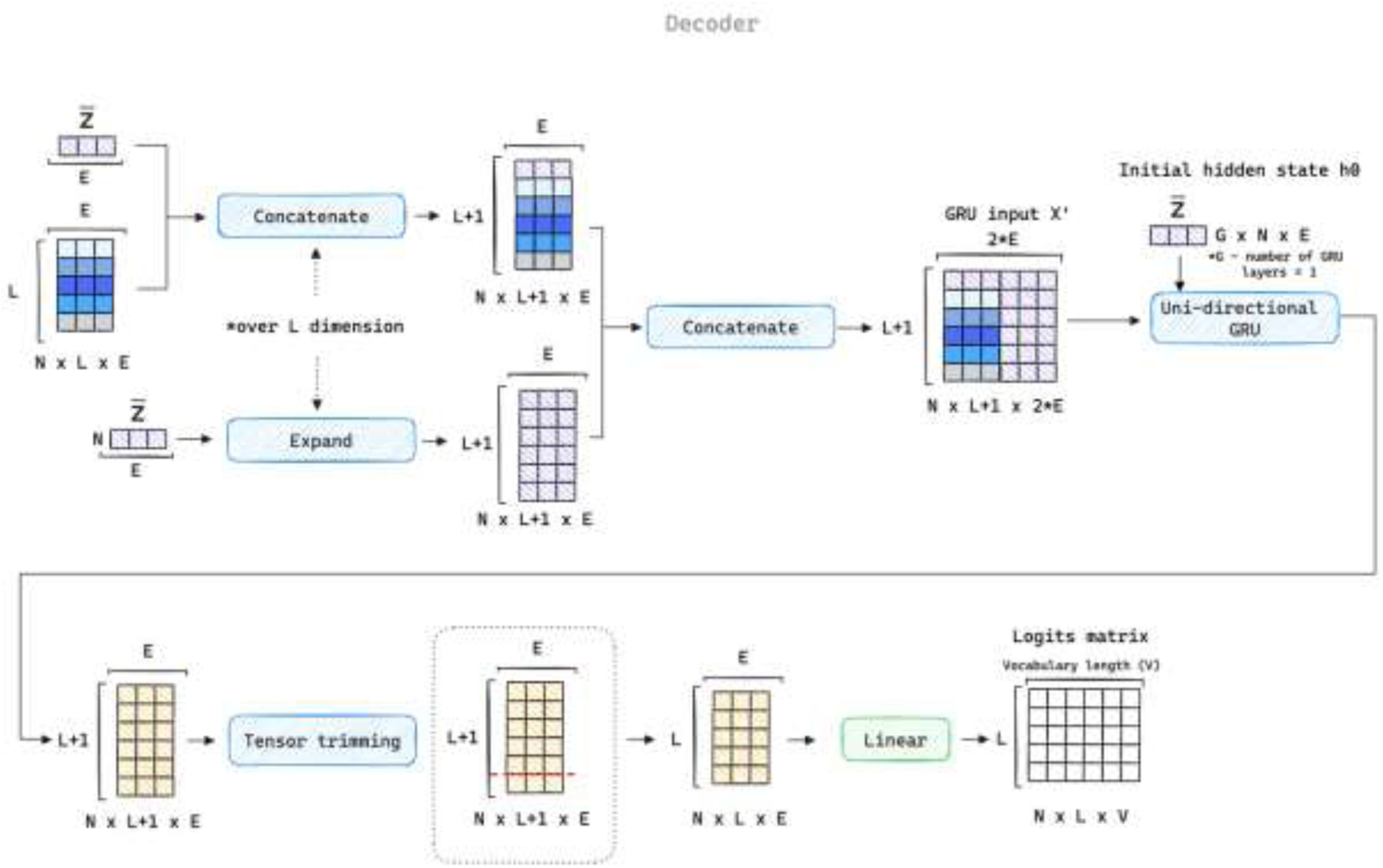
Scheme of the decoding process in WAE.

### Loss function

The loss function used herein is composed from the regularization, peptide length, and reconstruction losses as well as L1 (LASSO) and Kullback-Leibler (KL) encoder variance regularization terms, equations (1)-(2). The rationale for adding the L1 and KL variance regularizations is to avoid the variance collapse.^3,4^

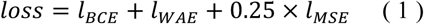

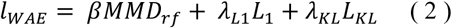

Where *l*_*BCE*_ is reconstruction loss, *l*_*WAE*_ is regularization loss, *l*_*MSE*_ is peptide length loss, *MMD*_*rf*_ is Maximum Mean Discrepancy (MMD) divergence metric with random features Gaussian kernel approximation, *β* is regularization coefficient, *L*_1_ is L1 penalty on log-variances, *λ*_*L*1_ is L1 regularization coefficient, *L*_*KL*_ is KL penalty on log-variances, *λ*_*KL*_ is KL regularization coefficient.

The WAE encoder seeks to force the aggregated encoded training distribution of peptides to match the prior, while providing detailed enough latent vectors to the decoder for accurate reconstruction.^5^ In more detail, the regularization penalty forces the aggregated posterior, i.e., the continuous mixture of all individual peptides’ distributions, to match the known prior distribution, i.e., Gaussian (normal). Herein, this is done as introduced by Gretton et al.^6^ and implemented by Das et al.^3^, through minimization of the Maximum Mean Discrepancy (MMD) divergence measure with random features Gaussian kernel approximation^7^ computed between the *z̅* vectors drawn from aggregated posterior and prior distributions in a given batch. The MMD is computed as described in the following steps: mapping of vectors drawn from aggregated posterior and prior to a common space with *rf*_*dim* dimensions through random features approximation of the Gaussian kernel, equation (3); calculation of the mean vector *μ*_*rf*_ of the transformed vectors ***Z***_***rf***_ in the random feature space, equation (4); computation of the MMD loss as the squared Euclidean distance between the mean vectors from the random feature space that allows to compare aggregated posterior and prior distributions, equations (5)-(6).

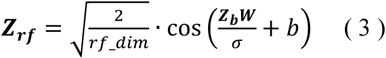

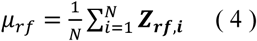

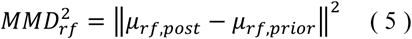

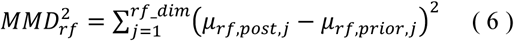

where *Z*_*b*_ is matrix of *z̅* vectors in a batch with dimensions (*N*, *E*), with *N* being the batch size and *W* being the random weight matrix with dimensions (*E*, *rf*_*dim*), *rf*_*dim* is the dimensionality of the random feature space, *b* is the vector of random bias value of length *rf*_*dim*, *σ* is the Gaussian kernel scale parameter; ***Z***_***rf***_ is a matrix of transformed vectors of dimensionality (*N*, *rf*_*dim*), *μ*_*rf*_ is the mean vector of transformed vectors ***Z***_***rf***_, *MMD*_*rf*_ is Maximum Mean Discrepancy metric for probability distributions comparison, *μ*_*rf*,*post*_ is a mean vector of the samples (vectors) drawn from the aggregated posterior distribution and mapped to random feature space, *μ*_*rf*,*prior*_ is a mean vector of the samples (vectors) drawn from the prior distribution and mapped to random feature space. The reconstruction cost function is the binary cross-entropy, equation (7). Since the peptides of the same length as the training set ones were prioritized, the length loss defined as MSE and multiplied by coefficient of 0.25 was added as an additional cost function, equation (8).

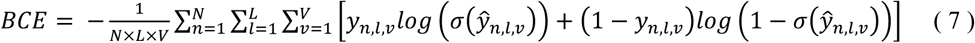

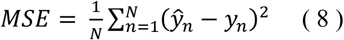

where *N* is the batch size, *L* is the maximum sequence length, *V* is the vocabulary size, *y*_*n*,*l*,*v*_ is the ground truth label for the v^th^ AA at the l^th^ position in the n^th^ sequence in one-hot encoded format, *y*^_*n*,*l*,*v*_ is the predicted unnormalized probability for the presence of the v^th^ AA at the l^th^ position in the n^th^ sequence, *σ* is the sigmoid function; *y*^_*n*_ is the predicted length of the n^th^ sequence, *y*_*n*_ is the true length of the n^th^ sequence.

### WAE and GTM parameters

In the WAE training, the AdaBelief^8^ optimizer was used with a learning rate of 0.0005 and dropout rate of 0.2. The loss coefficients were set to 0.01 for *β* and 0.001 for both *λ*_*L*1_ and *λ*_*KL*_. The stochastic weight averaging (SWA) was applied for enhanced generalization. The GTM β parameter optimized by the EM algorithm was determined to be 0.04015. Optimal predictive performance of class landscapes was observed with the manifold with the following hyperparameters: 37 nodes, 22 RBFs, an RBF width of 0.3, regularization coefficient of 0.087096. To increase the resolution, the map was resampled to a finer grid of 30 to 30 dimensions, overall comprising 900 nodes.

### Statistical metrics for evaluation of QSAR models

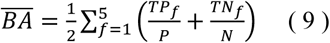

where *TP_f_* – the number of truly active peptides predicted as active in the fold f, *TN_f_* – the number of truly inactive peptides predicted as inactive in the fold f, N – the total number of peptides belonging to the inactive class, P – the total number of peptides belonging to the active class.

